# Mosquito infection-mediated trait variation alters temperature-dependent transmission of West Nile virus

**DOI:** 10.64898/2026.08.06.743329

**Authors:** RL Fay, M Cruz-Loya, EM Banker, EA Mordecai, AT Ciota

**Affiliations:** Arbovirus Laboratory, Wadsworth Center, New York State Department of Health, Slingerlands, NY, USA; Department of Biomedical Sciences, University at Albany College of Integrated Health Sciences, Albany, NY, USA; Biology Department, Stanford University, Stanford, CA, USA

**Keywords:** West Nile virus, Temperature, Thermal Performance Curve, R_0_

## Abstract

Rising global temperatures are reshaping species interactions and the ecological conditions governing vector-borne disease transmission. Although previous studies show that West Nile virus (WNV) infection alters mosquito longevity, fecundity, blood-feeding behavior, and the thermal performance of these traits, trait-based R₀ models largely rely on data from uninfected mosquitoes, implicitly assuming homogeneous vector populations. This overlooks infection-induced trait variation that may influence transmission dynamics. Here, we examined how temperature, infection status, and viral strain interact to shape transmission potential for WNV in *Culex pipiens*. Life-history traits of WNV-exposed and unexposed mosquitoes were measured across constant temperatures ranging from 10°C to 33°C, as well as under a fluctuating temperature regime of 25°C ± 5°C. These data were used to generate thermal performance curves and estimate temperature-dependent relative R₀ across treatments. Infection altered the thermal performance of mosquito life-history traits, vector competence, and overall transmission potential. We also found evidence for a bimodal effect of temperature on vector competence, potentially driven by tradeoffs between viral replication and mosquito immune responses. Incorporating infection-sensitive traits into relative R₀ calculations reduced estimated transmission intensity across much of the thermal range without shifting thermal optima or limits, suggesting that current models may overestimate transmission.

## Introduction

Arthropod-borne viruses (arboviruses) cause more than 140 recognized human diseases globally, with several of the most widespread pathogens belonging to the Orthoflavivirus genus, including West Nile virus (WNV). WNV is maintained and transmitted primarily by Culex mosquitoes, particularly Culex pipiens[1–3]. Following its introduction into the Western Hemisphere in 1999, WNV was associated with outbreaks of neuroinvasive disease and rapidly expanded across North America and subsequently throughout much of the Americas[4–6]. Over the past several decades, global temperatures have risen substantially, creating new challenges for predicting and managing mosquito-borne disease risk[7,8]. Because mosquitoes and the viruses they transmit are ectothermic, their physiology and performance are strongly influenced by environmental temperature[9,10]. Temperature affects numerous components of the transmission cycle, including mosquito survival, development, reproduction, biting behavior, and viral replication within the vector[11–13]. As a result, changes in thermal conditions can alter both the intensity and duration of WNV transmission[8,13].

The basic reproduction number (R_0_) is used to predict the transmissibility of a pathogen, and a modified Ross-MacDonald R_0_ was developed to account for temperature-dependent life history traits for mosquito-borne pathogens[11–15]. Thus far, such models have utilized empirical data to generate thermal performance curves (TPCs)[11–13,16]. Previous studies have shown that exposure to WNV can influence how mosquito life-history traits respond to temperature, including adult lifespan, blood-feeding, and oviposition[17–19], yet current R_0_ models utilize non-infectious mosquito life history trait data to predict the influence of temperature on vector-borne diseases. However, vector-borne disease transmission requires a combination of traits in unexposed vectors—including population dynamics and blood feeding on potentially infectious hosts—and virus-exposed vectors—including survival through the extrinsic incubation period and subsequent blood feeding on potentially susceptible hosts. If virus-exposed mosquitoes have systematically different thermal responses in key life history traits relative to unexposed mosquitoes, models that ignore this heterogeneity may fail to capture the true magnitude and thermal performance of transmission. Yet, the combined effects of WNV exposure and temperature on mosquito life history traits and transmission have yet to be investigated.

WNV strains have evolved under rising temperatures, potentially influencing mosquito life-history traits and transmission dynamics of WNV vectors[16]. These evolutionary changes are reflected in shifts in dominant viral genotypes over the last 25 years from the original NY99 genotype to the WN02 genotype, which emerged in 2002 and rapidly spread across the US[20]. Later, the NY10 genotype became dominant in New York State (NYS) following its emergence in 2010, and subsequently spread throughout the U.S.[21,22]. Canonical estimates of WNV transmission using modified Ross-MacDonald R₀ models have relied primarily on data from WN02 infections[11,12], which may overlook the ecological consequences of viral evolution. Prior research indicates that NY10 genotype strains, having evolved under higher temperature regimes, exhibit increased transmissibility by *Cx. pipiens* at elevated temperatures[16], underscoring the importance of considering virus–vector–environment interactions when evaluating disease risk.

Thus, in this study we address a two-fold knowledge gap: understanding the impact of virus exposure on the thermal responses of mosquito life history and transmission, and how this may differ among virus genotypes with different evolutionary histories. We experimentally measured adult mosquito life history traits across temperatures and infection status for both WN02 and NY10 genotype strains to obtain more accurate predictions of WNV risk. These data were utilized to generate TPCs and incorporated into a modified relative R_0_ model that accounts for infection-dependent mosquito life history traits, which we compare to the standard approach that uses solely TPCs measured in unexposed mosquitoes. Our approach predicts reduced WNV risk compared to models that assume mosquito traits do not vary with infection status due to a decrease in biting rates and lifespan in WNV-infected mosquitoes, which is consistent across both viral strains. Together, these results demonstrate that more accurately predicting the risk of vector-borne diseases under climate change requires considering how the interaction of temperature and infection influence vector biology.

## Methods

### Viruses

The two strains used in this study were collected from NYS mosquito surveillance and genetically analyzed using protocols outlined in prior studies[16,21]. The 2003.2 strain used was isolated in 2003 (DQ164189) from an American crow (*Corvus brachyrhynchos*) found in Albany County, NY, which was initially amplified on Vero cells for sequencing, and then later amplified on C6/36 cells (*Aedes albopictus*, ATCC, Manassas, VA) for downstream use. The 2017.1 strain (MT968008.1) utilized in this study was isolated from *Culex* mosquito surveillance pools in 2017. The 2017.1 strain was amplified one time on Vero and C6/36 cells. After 5 days of amplification on C6/36 tissue culture supernatant was harvested and stored in 20% FBS at -70°C. The more contemporary strain of WNV, 2017.1, has 8 amino acid substitutions that are not shared with the historic 2003.2 strain[16].

### Mosquito rearing

*Cx. pipiens* colony mosquitoes were originally collected in Pennsylvania in 2004 (courtesy of M. Hutchinson) and have been highly colonized at the Wadsworth Center Arbovirus laboratory. *Cx. pipiens* mosquitoes were maintained in 30.5-cm^3^ cages in an environmental chamber at 27 ± 2°C with a relative humidity of 45-65% and a photoperiod of 16:8 (L:D) h and provided cotton pads with 10% sucrose *ad libitum*. Adults were grouped and housed in gallon-size cardboard containers via aspirator as they emerged and held for 1 day to allow mating.

### Blood feeding and experimental infection

Four to seven-day-old adult females were collected and fed on doses of WNV of 6.92 log_10_ PFU/mL for the 2003.2 strain or 7.38 log_10_ PFU/mL for the 2017.1 strain. Blood meals consisted of a 4:1 mixture of diluted virus stock and chicken blood (Colorado Serum Company, Denver, CO), and a final concentration of 2.5% sucrose. Additionally, a non-infectious blood meal was also provided to separate groups of mosquitoes to allow the comparison of WNV-exposed and unexposed life history traits. Following one hour of feeding using an artificial feeding chamber (Hemotek, Blackburn, UK) at 37°C, mosquitoes were anesthetized, and the engorged females were collected, knocked down using CO_2_, and placed in individual 50 mL conical tubes placed in Styrofoam rack, with a hole in the top of the conical covered in mesh, a hole in bottom of the conical to allow addition of water for laying, and a dental dam around bottom of the conical to hold the tube in place. Cotton pads soaked in 10% sucrose ad libitum were placed on top of the conicals. Individual conicals were then held at various temperatures, including 10°C, 15°C, 20°C, 25°C, 30°C, and 33°C at constant temperatures, and additionally at a cycling temperature between 20-30°C, around a mean of 25°C. The sample size was 45-50 WNV blood-fed mosquitoes for each strain and temperature, and 30 non-infectious blood-fed mosquitoes per temperature.

### Mosquito Survival, Blood Feeding, and Fecundity

Survival was checked daily for all treatment and temperature groups. The bottom of the conicals were checked daily for eggs; when eggs were observed in a conical, the egg raft was removed using a wooden stick. Additional water was added as needed to the bottom of the conicals. To measure blood-feeding rates, mosquitoes were starved overnight for 12-24 h and offered 200-μL defibrinated chicken blood (Colorado Serum Company, Denver, CO) with 2.5% sucrose via absorbent pad for 2-h 1x per week and evaluated for feeding activity after 2 hours.

### Vector competence

Upon death, the individual mosquito’s legs and body were saved separately with a 4.5 mm zinc-plated steel ball (BB) (Daisy, Dallas, TX) in 500 μL mosquito diluent (MD; PBS with 20% FBS, 100 μg/mL penicillin/streptomycin, 10 μg/mL gentamicin, 1 μg/mL amphotericin B) at -80°C. To determine positivity, thawed samples were homogenized at 24Hz for 3 minutes, followed by centrifugation at 1200 rpm for 3 minutes, and RNA was extracted using a MagMAX-96 Viral RNA Isolation Kit (Thermo Fisher, Waltham, MA, USA) on a MagMax Express-96 Magnetic Particle Processor (Applied Biosystems, Waltham, MA, USA). Real-time quantitative RT-PCR was completed using qScript One-Step RTqPCR ToughMix Low Rox (QuantaBio, Beverly, MA, USA) and analyzed on Quant Studio 5 (Thermo Fisher, Waltham, MA, USA). WNV primers and probes were designed as previously described and copy standards were utilized for quantification[23]. Vector competence was assessed by quantifying the proportions of infected (positive bodies) and disseminated (positive legs) mosquitoes for 45-50 mosquitoes per virus strain. Strain-specific data were analyzed and compared using GraphPad Prism 9.

### Thermal performance curves

A Bayesian approach was used to estimate TPCs for mosquito lifespan, biting rate, vector competence and oviposition (see Appendix for model fitting details). Separate curves were fit to mosquitoes unexposed to WNV and, for each virus strain, to exposed mosquitoes that did not develop a detectable infection, and to mosquitoes that became infected. In this analysis, mosquitoes were considered infected if they tested positive for WNV in either the legs or bodies (i.e., pooling the infected and disseminated populations). Traits with a unimodal response to temperature (lifespan, biting rate and oviposition) were modeled with a flexTPC functional form[24], which allows comparing curves of different shapes with a single model.

Unexpectedly, vector competence exhibited bimodal, rather than unimodal, variation with temperature. For both WNV strains, vector competence decreased at 15°C relative to 10°C, increasing again at higher temperatures. To describe the thermal performance of this trait, we developed a novel semi-mechanistic model where vector competence is assumed to result from a tradeoff in temperature sensitivities between the viral replication rate and mosquito immunity, leading to a bimodal vector competence curve (see Appendix for details). A Bayesian approach was used to fit this model to the infection and dissemination data, inferring the underlying viral replication rate and mosquito immunity curves to generate an overall bimodal thermal performance curve for these vector competence traits.

Models were fit using Markov Chain Monte Carlo (MCMC) with the r2jags R package, an interface for the JAGS (Just Another Gibbs Sampler) program[25]. For all traits except vector competence, eight independent MCMC chains were run for 300,000 iterations, discarding the first 50,000 iterations as burn-in. The resulting MCMC chains were thinned, saving every eight iterations. As the vector competence model exhibited slower convergence, eight independent MCMC chains were run for 2,080,000 iterations, discarding the first 80,000 iterations as burn-in and thinning the chains by saving every 64 iterations. Chain convergence was monitored both by visual inspection of trace plots and density plots of the individual chains and by ensuring the potential scale reduction factor *R̂* < 1.01 and *n_eff_* > 10000 for all parameters. A summary of the posterior distribution (mean, 95% credible interval [CI], *R̂*, *n_eff_*) for all model parameters corresponding to the TPC fit for each trait is available in the Appendix.

### Temperature-dependent Relative R_0_

The temperature-dependence of WNV transmission was evaluated using a relative R_0_ approach^12^ that combines the TPCs estimated here with previously published estimates of the thermal performance of other *Cx. pipiens* life-history traits and the WNV pathogen development rate (see Appendix for details). Standard approaches to calculate relative R_0_ implicitly assume that mosquito life-history traits do not change in infected mosquitoes. To account for infection-dependent changes, we developed a modified relative R_0_ model that incorporates infection-dependent biting rates and lifespan (Equation 1, see Appendix for model derivation and details).

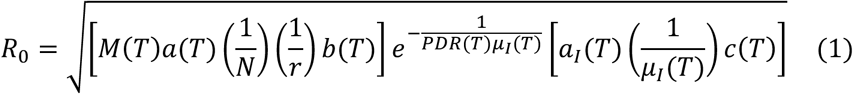

where *M*(*T*) is mosquito abundance, *a*(*T*), *a_I_*(*T*) the biting rates for unexposed and infected mosquitoes, respectively, *N* the number of mosquitoes, *r* the recovery rate, *b*(*T*) the probability of a mosquito getting infected when biting an infected host, *c*(*T*) the probability of a host getting infected when bitten by an infected mosquito, *PDR*(*T*) the pathogen development rate, and *μ*(*T*), *μ_I_*(*T*) are the mortality rates (inverse of lifespan) for unexposed and infected mosquitoes. As in previous work, we ignore *N* and *r*, which are temperature-independent traits that are difficult to estimate. This yields a relative *R*_0_ that is proportional to *R*_0_ and has the same temperature-dependence, but which has an arbitrary scale that cannot be directly interpreted in terms of the number of secondary cases arising from an infection:

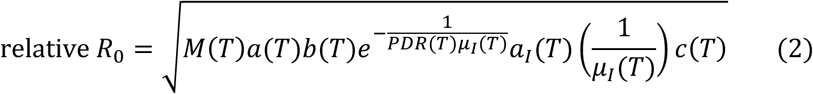

These expressions generalize previous models for *R*_0_ and relative *R*_0_ based exclusively on traits of uninfected mosquitoes, which are a special case of the expressions shown here when the mortality and biting rates of unexposed and infected mosquitoes are equal (i.e. when *μ*(*T*) = *μ_I_*(*T*) and *a*(*T*) = *a_I_*(*T*)). As in previous work, we estimate the mosquito abundance *M*(*T*) from various mosquito life-history traits[12] (see Appendix). We ignore the effects of infection in oviposition when estimating mosquito abundance, as these effects are subtle and the proportion of infected mosquitoes in the field is typically low, making mosquito abundance primarily determined by the traits of uninfected mosquitoes.

When estimating relative *R*_0_ with either equation, we use the TPCs for lifespan, biting rates, and oviposition estimated in this work, including viral strain-specific estimates for *a_I_*(*T*), *μ_I_*(*T*) and vector competence *bc*(*T*), which is the product of *b*(*T*) and *c*(*T*) in the expressions above. For traits not measured here, we used previously estimated TPCs for life history traits in *Cx. pipiens* from Shocket *et al* 2020 (see Appendix for details).

## Results

### Infection status influences mosquito life history traits

*Cx. pipiens* mosquitoes were fed either an infectious bloodmeal of 2003.2 or 2017.1 WNV strains or a non-infectious bloodmeal. Fully engorged females were individually housed at 10, 15, 20, 25, 30, or 33°C as well as an additional treatment cycling from 20-30°C with a mean of 25°C. Mosquito life history traits were recorded across temperatures and utilized to generate TPCs (Fig. 1, S1, S5-7, and Table S1-3). We find that mosquito life history traits vary between unexposed and WNV-infected mosquitoes and between WNV strains (Fig. 1 and Fig. S5-7).

**Fig 1.**
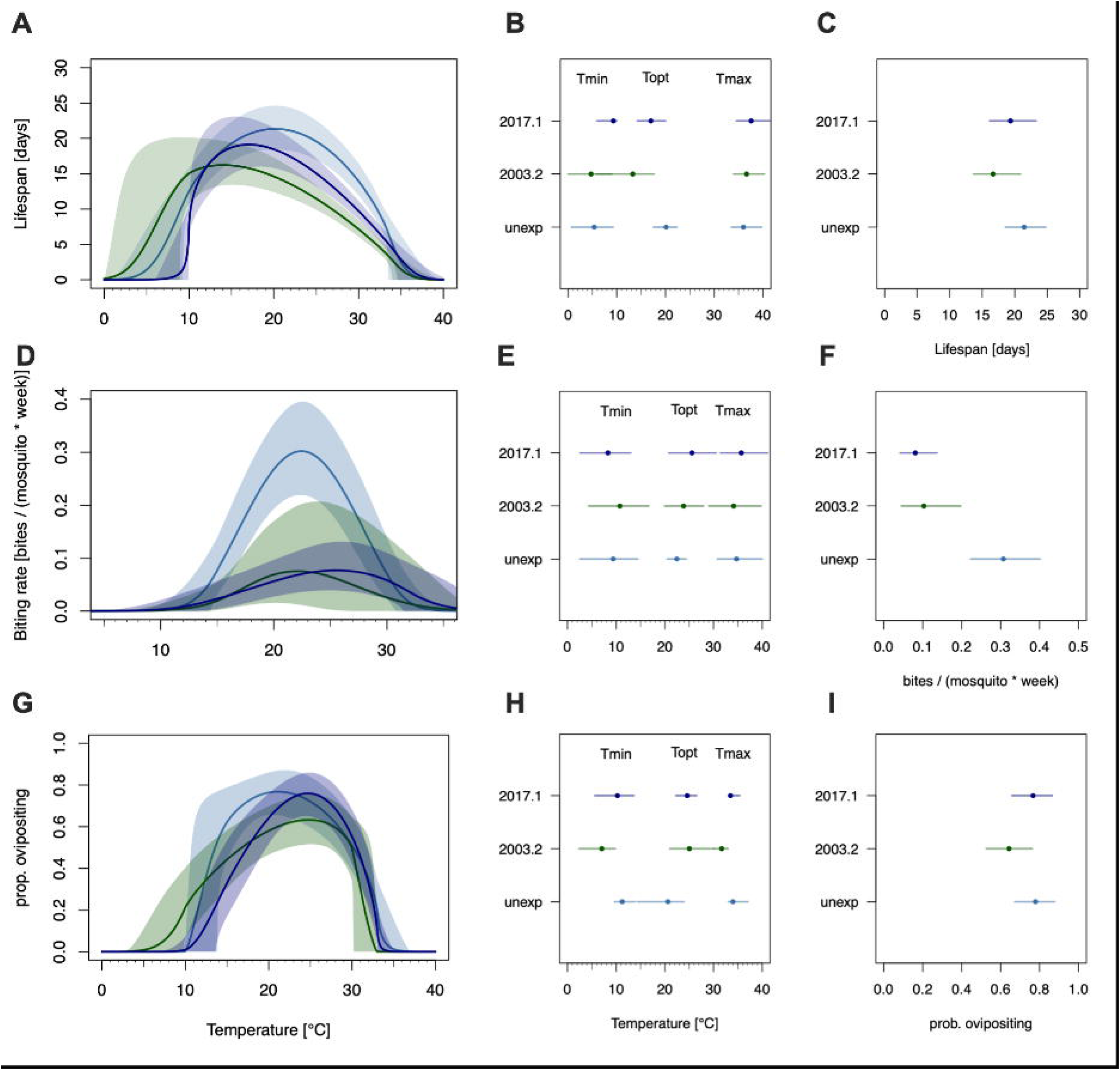
Thermal performance curves of *Cx. pipiens* life history traits. **A,D,G)** Inferred thermal performance curves of life history traits (**A**: adult lifespan, **D**: biting rate, **G**: oviposition) of *Culex pipiens* mosquitoes of varying infection status (blue: unexposed, green: infected with 2003.2 strain, purple: infected with 2017.1 strain). Solid lines are posterior distribution means; shaded areas are 95% credible intervals. Data points not shown here for clarity, but individual curve fits are shown in Fig S6-S8). **B,E,H)** Estimated thermal minimum, optimum, and maximum of *Culex pipiens* life history traits (**B**: adult lifespan, **E**: biting rate, **H**: oviposition). Points correspond to posterior means and lines to 95% credible intervals. **C, F, I)** Estimated peak rate for each life history trait (**C**: adult lifespan, **F**: biting rate, **I**: oviposition).

Across temperatures, virus exposure significantly affected the thermal response of adult mosquito lifespan (Fig. S1 and Table S1; *p* ≤ 0.05 two-way ANOVA with Tukey pairwise-comparison). At 10°C, strong genotype-specific survival differences were evident, with 2003.2-exposed uninfected mosquitoes living longer than several 2017.1 groups. No significant differences were detected at 15°C. Effects were limited at 20–25°C, but significant survival costs of virus exposure reemerged at 30–33°C, particularly among infected groups. At the cycling treatment, mosquitoes infected with the strain 2017.1 had significantly reduced lifespan compared to unexposed and 2003.2 infected mosquitoes (Fig. S8A; p ≤ 0.05, one-way ANOVA with multiple comparisons, Tukey’s post-test). Our TPC results demonstrate clear evidence of decreased lifespan in WNV-infected mosquitoes relative to unexposed mosquitoes across a temperature range of 15-34°C for the 2003.2 strain and a narrower range of 20-33°C for the 2017.1 strain (Fig. 1A and S4-5). The estimated TPCs for lifespan in WNV-exposed mosquitoes (regardless of infection status) also have an apparent lower thermal optimum than for unexposed mosquitoes, although this finding is only conclusive for 2003.2 exposed uninfected mosquitoes, as other conditions have overlapping credible intervals (Figs. 1B and S5).

Blood-feeding was found to be strongly temperature-dependent (Fig. S1 and Table S2; *p* ≤ 0.05 Fisher’s exact test with Bonferroni correction). Mosquitoes did not blood feed at extreme temperatures (10°C and 33°C), with minimal feeding at cool temperatures (15°C), and peak feeding at moderate to warm temperatures (20–28°C), especially at cycling temperature (Fig. S1B). Among exposed mosquitoes, infection and dissemination status have only minor effects, with modest increases or decreases in feeding at specific temperatures though not significant, but overall temperature and virus exposure were the dominant drivers of feeding behavior (Figs. 1, S1B, and Table S2). Our TPC results show a substantially reduced biting rate in WNV infected mosquitoes (peak biting rate of 0.103 95%CI:[0.045, 0.196] for the 2003.2 strain and 0.081 95%CI:[0.042, 0.136] for the 2017.1 strain) relative to unexposed mosquitoes (0.307 95%CI:[0.223, 0.401]): 66% and 74% reductions in peak biting rate, respectively (Fig. 1D). Interestingly, exposed uninfected mosquitoes also had decreased biting rates relative to unexposed mosquitoes, although to a lesser extent and with more individual variability than infected mosquitoes (Fig. S6).

Oviposition varied by temperature and infection status, with exposed uninfected mosquitoes consistently having reduced oviposition across temperatures and virus strains (Fig. S1C and Table S3; *p* ≤ 0.05 Chi-squared with Monte Carlo and Bonferroni correction). At lower temperatures (10°C), differences are minimal across groups. As temperatures rise (15–30°C), significant differences emerge, particularly between unexposed mosquitoes and both 2003.2 and 2017.1 exposed mosquitoes, and across strain and infection status (Fig. S1C and Table S3). At the fluctuating temperature, mosquitoes infected with the 2003.2 strain had similar proportions ovipositing compared to unexposed mosquitoes, but 2017.1 infected as well as virus-exposed uninfected mosquitoes had a lower proportion ovipositing (Fig. S8C; p ≤ 0.05, one-way ANOVA with multiple comparisons, Tukey’s post-test). The TPC results show that exposure to WNV leads to decreased oviposition rates at low temperatures (∼10-20°C) and an apparent increase to the oviposition thermal optimum, with point estimates around 25-28°C compared to 20.6°C 95%CI:[14.3°C, 23.9°C] for unexposed mosquitoes (Fig. 1). However, this finding is only conclusive for 2017.1 exposed uninfected mosquitoes (28.2°C 95%CI:[24.2°C, 31.7°C]), as in other cases the credible intervals overlap (Fig. 1 and S7).

### Vector competence of WNV-exposed Cx. pipiens

To assess strain-specific differences in vector competence across temperature, we collected the bodies and legs of females upon death to determine infection and dissemination, respectively. The overall trend in vector competence data shows that as temperature increases, *Cx. pipiens* vector competence also increases, with a notable exception at 10°C described below. Significant differences in vector competence by *Cx. pipiens* were found at the fluctuating temperature wherein the number of females with disseminated 2003.2 infections was higher than the number of mosquitoes with a disseminated 2017.1 infection (Figs. 2, S8E, and Table 1; *p* ≤ 0.05 Chi-squared and p ≤ 0.05, one-way ANOVA with multiple comparisons, Tukey’s post-test). At the higher temperatures of 30°C and 33°C, *Cx. pipiens* showed significantly increased vector competence of the 2017.1 strain compared to the 2003.2 strain (Fig. 2 and Table 1; *p* ≤ 0.05 Chi-squared). Interestingly, at 10°C we saw an increase in dissemination across both strains compared to that at 15°C (Fig. 2 and Table 1).

**Fig 2.**
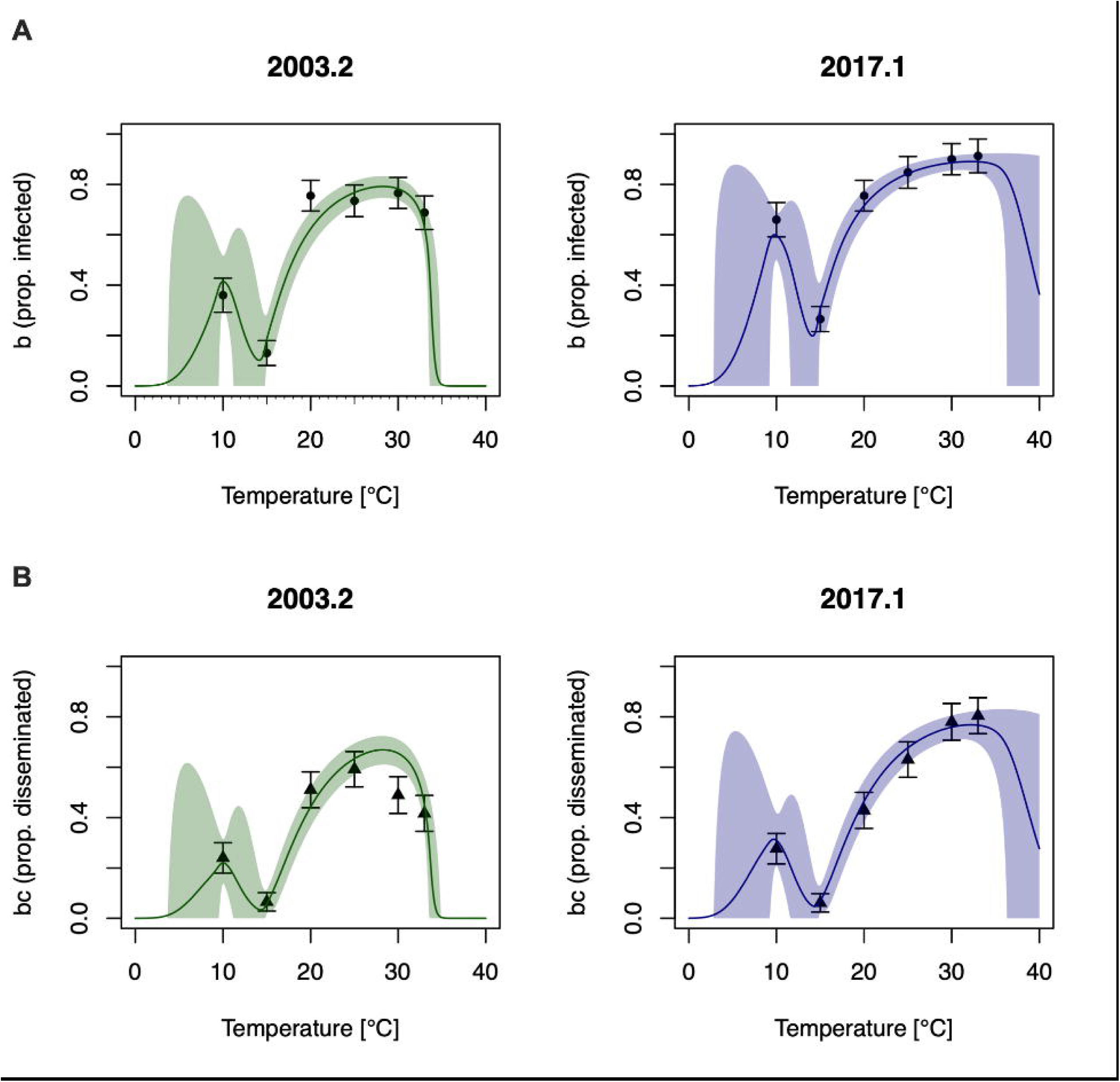
Vector competence has a complex, non-unimodal relationship with temperature. **A)** Observed proportion infected are shown as dots and **B)** proportion disseminated shown in triangles with standard errors shown as error bars. Lines represent posterior means and shaded regions 95% credible intervals of a semi-mechanistic model assuming vector competence arises from a tradeoff between viral replication and mosquito immunity (see Methods and Appendix).

**Table 1.**
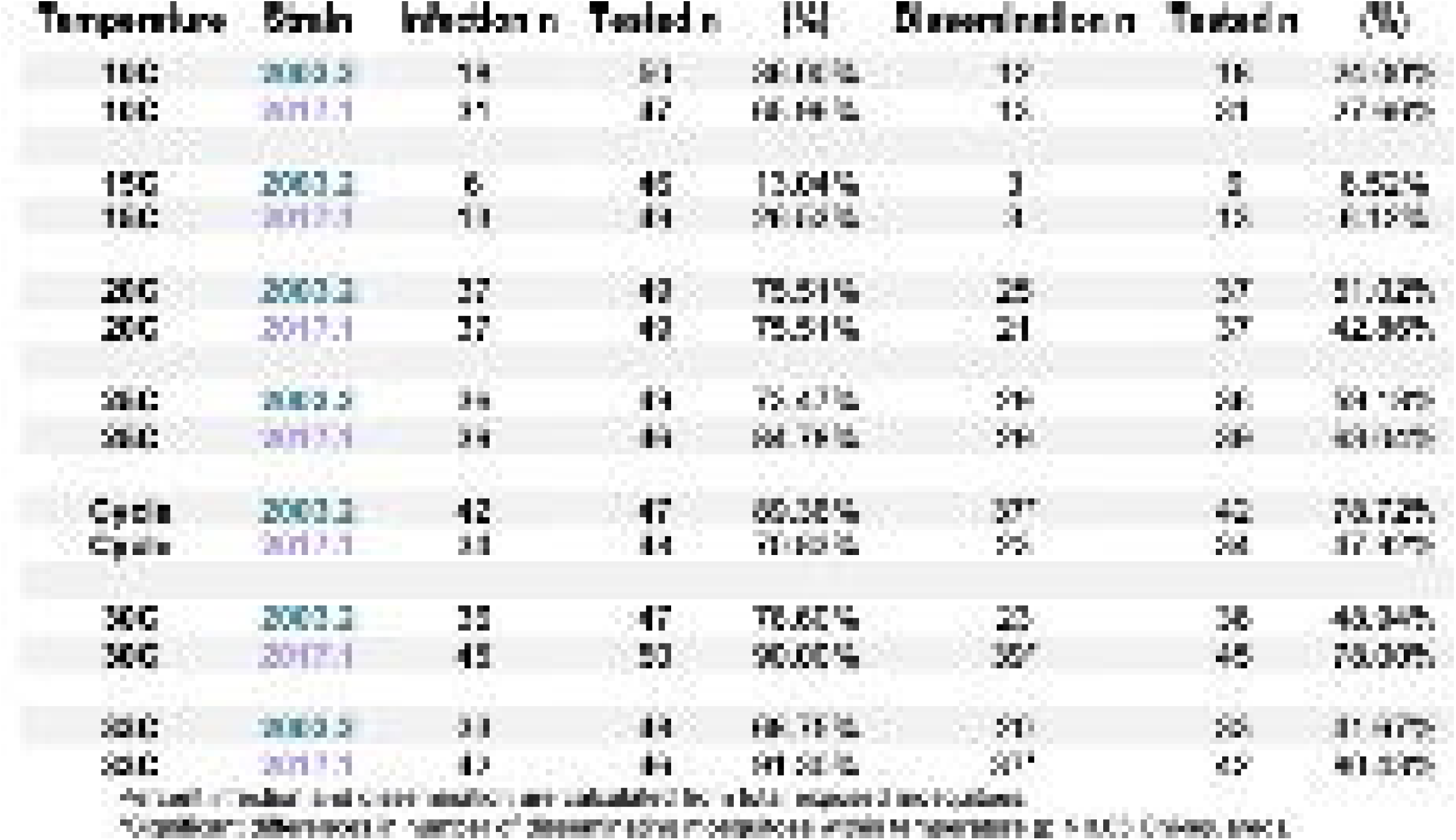
*Culex pipiens* infection rates and dissemination rates after exposure to 2003.2 and 2017.1 West Nile virus strains.

For both the 2003.2 and 2017.1 strains, the proportion of infected (*b*) and disseminated (*bc*) mosquitoes exhibited a non-unimodal relationship with temperature, with an initial decrease between 10°C and 15°C followed by an increase to a peak at higher temperatures (Fig. 2). To describe this relationship, we developed a semi-mechanistic model based on the hypothesis that the temperature sensitivity of vector competence arises from a tradeoff between the thermal responses of viral replication and mosquito immunity, which are represented as latent (unobserved) TPCs inferred from the vector competence data (see Fig. S3 and Appendix). This model predicts a bimodal TPC for vector competence wherein low mosquito immunological defense allows for moderate vector competence at 10°C, followed by a decrease at 15°C as immunological defense increases, followed by an increase at 20°C and above as rapid viral replication outpaces the mosquito immune defense. This model fits the data well (Fig. 2), showing that this hypothesis is a plausible explanation for the observed non-unimodal relationships. However, it should be considered as tentative, as experimental confirmation is needed to validate the role of mosquito immunity in shaping the vector competence TPC.

### Temperature-dependent Relative R_0_

Current approaches to model WNV transmission implicitly assume that mosquito life-history traits are identical for unexposed and infected mosquitoes. Our experimental results show clear changes in biting rates and lifespan in infected mosquitoes, suggesting that transmission models need to be revised to incorporate these trait differences. To evaluate the effect of incorporating infection-varying traits in WNV transmission, we developed a modified temperature-dependent relative R_0_ model that incorporates traits of both unexposed and infected mosquitoes (as appropriate in each stage of the transmission cycle) and compared its predictions with an existing approach using only traits from unexposed mosquitoes (Fig. 3, see Appendix for derivation). Our revised transmission model predicts substantially lower overall WNV transmission due to decreased biting rates and lifespan in infected mosquitoes but does not meaningfully change the predicted thermal optimum or limits for transmission (see Appendix). Additionally, we do not find evidence for strain-specific differences between the 2003.2 and the 2017.2 relative R_0_ curves.

**Fig 3.**
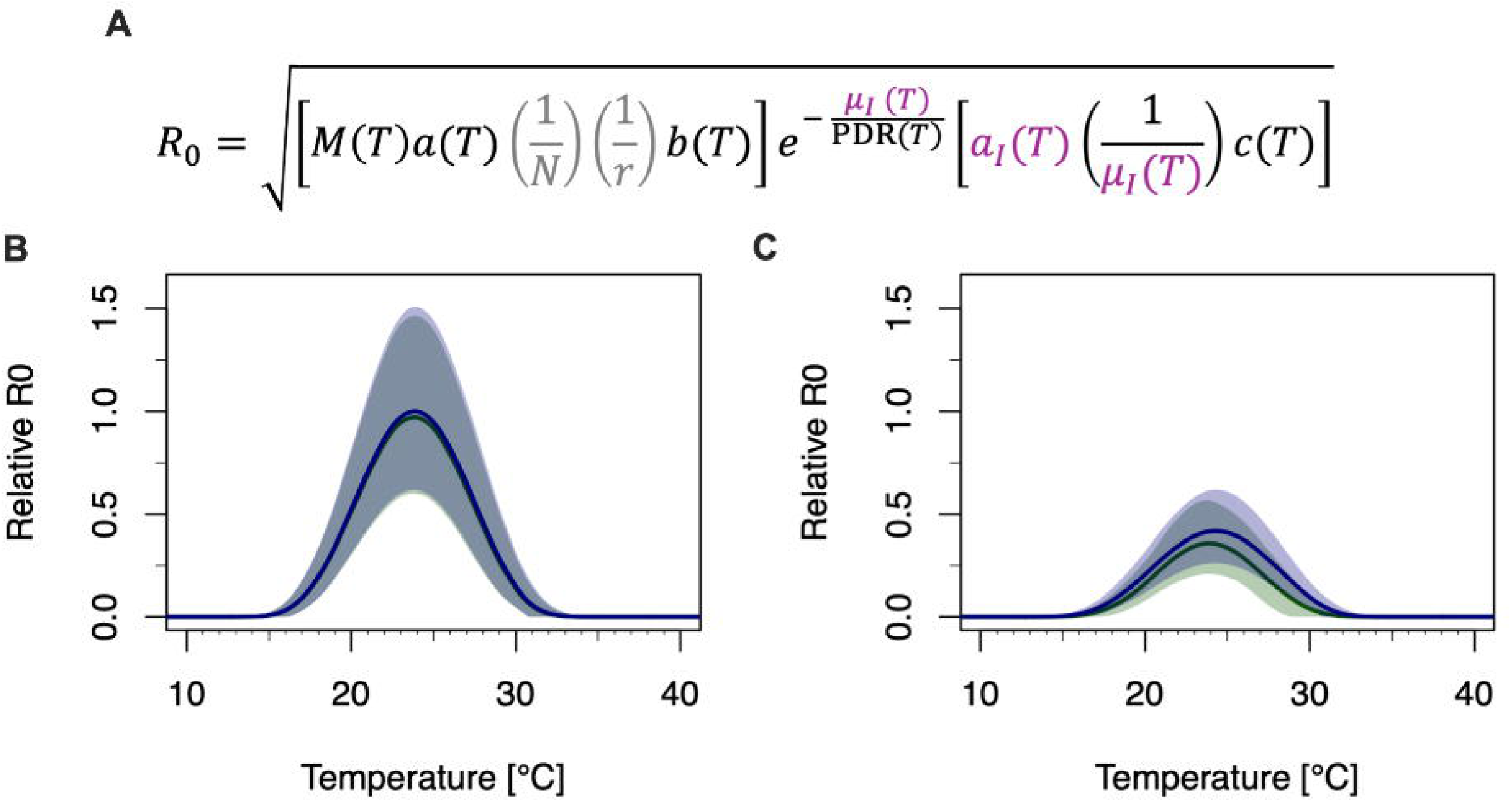
Incorporating infection-sensitive mosquito life history traits modifies WNV transmission estimates. **A)** The basic reproductive number (R_0_) for WNV transmission can be broken down into three factors that depend on various temperature-dependent mosquito and viral traits: i) the number of mosquitoes that become infected from biting a single infected host, ii) the proportion of infected mosquitoes surviving the extrinsic incubation period (EIP) and iii) the number of hosts that become infected from the surviving infected mosquitoes (see Methods and Appendix for notation used for each trait). **B**) standard approach to calculate assuming life history traits do not change with exposure status (which only uses traits of unexposed mosquitoes *a*(*T*), *μ*(*T*)). **C**) Novel transmission model that incorporates infection-dependent changes in mosquito biting rate *a_I_*(*T*) and mortality rate *μ_I_*(*T*) of infected mosquitoes in factors ii and iii (modified traits are shown in purple in formula). Following previous work^11^, we calculate relative R_0_ by ignoring temperature-independent traits that are difficult to estimate (shown in gray). Relative R_0_ is proportional to R_0_ (and has the same temperature-dependence) but has an arbitrary y-axis scale. As such, these estimates are informative about relative changes in transmission but cannot be interpreted as the number of secondary cases.

## Discussion

Our results demonstrate that WNV exposure and infection have meaningful effects on adult *Cx. pipiens* life-history traits and their thermal responses, reducing overall transmission potential while thermal optima and limits remained broadly similar across treatments (Fig. 3). The impacts of WNV exposure on traits such as lifespan, biting, and oviposition varied across temperatures (Figs. 1, S1, Tables S1–S3), highlighting the importance of environmental context in shaping host–pathogen interactions. Our TPC analysis found the most pronounced effect of WNV infection on biting behavior, emphasizing that vector competence is not static but dynamically influenced by both viral and environmental factors. Notably, we observed an unexpected increase in *Cx. pipiens* competence for WNV at 10°C, a pattern not previously reported in the literature but consistent across both virus strains and all experimental replicates, as well as increased vector competence of the contemporary WNV strain (2017.1) compared to the historic (2003.2) strain at high temperatures (Fig. 2 and Table 1). Although the changes in TPC shape for exposed mosquitoes did not meaningfully alter the R₀(T) curve in this system, similar changes in other mosquito–pathogen systems could shift the thermal minimum, optimum, or maximum for transmission. Generating empirically grounded, ecologically relevant trait data across realistic temperature gradients will be critical for accurately predicting WNV transmission in a warming world.

Life-history traits of *Cx. pipiens* varied across temperature, infection status, and viral strain, highlighting the dynamic interplay between mosquito physiology, behavior, and environmental conditions. The clearest difference between unexposed and WNV-exposed mosquitoes was in blood-feeding rate, with a higher proportion of unexposed individuals successfully taking a blood meal compared to mosquitoes with disseminated infections (Fig. S10). Among the two virus strains, important differences in competence at high temperatures likely reflect the combined effects of intrinsic viral phenotype and host behavioral ecology[19,26,27]. Although our experimental design required a blood meal for inclusion, and few exposed mosquitoes took multiple sequential meals, feeding frequency still varied across temperatures and between strains (Fig. S2). Because blood feeding directly governs transmission potential in natural populations, such strain- and temperature-dependent differences in feeding behavior merit further ecological investigation[26–28].

Mosquitoes maintained under realistic, fluctuating temperature regimes showed vector competence patterns that diverged from those observed under static 25°C conditions (Table 1 and Fig. S8DE). At the cycling temperature (mean ∼25°C), strain 2003.2 exhibited significantly higher dissemination than 2017.1, whereas under static conditions, 2017.1 showed higher dissemination that was not statistically significant. These differences underscore the ecological importance of incorporating diurnal and seasonal temperature variation into laboratory and field studies[29–32]. Together, our findings demonstrate that temperature regime, infection status, and viral strain interact to shape mosquito life history and transmission potential, emphasizing the need for empirically derived, infection-sensitive trait data to improve predictions of vector-borne disease risk with changing climates. Our study focused on *Cx. pipiens*, leaving open whether similar patterns occur in other key WNV vectors (e.g., *Cx. quinquefasciatus, Cx. tarsalis*), across additional viral strains, or in other mosquito–arbovirus systems. Therefore, pathogen-induced modification of mosquito traits may be a largely overlooked contributor to ecological variability in mosquito-borne transmission systems and deserves expanded investigation.

Although the fitness costs of WNV infection in Cx. pipiens have been reported previously, these effects have not been evaluated across ecologically relevant thermal gradients[17]. Prior studies documented infection-associated differences in survival, biting, and oviposition survival [17–19], but whether these costs vary across temperature remained unknown. Our results demonstrate that these fitness and behavioral effects persist across a range of temperatures, highlighting their potential influence on host contact rates, reproductive output, and ultimately transmission potential. Importantly, these effects were not strain-specific, suggesting that infection-mediated trait modification may reflect a generalized physiological cost of infection rather than lineage-specific interactions. This interpretation is consistent with broader evidence from other mosquito–arbovirus systems, including studies of dengue virus and Zika virus, where infection has been shown to alter feeding behavior, fecundity, and survival [17,33].

Another novel finding of this study was a non-unimodal pattern in vector competence across temperature. This study represents the first assessment of WNV dissemination in *Cx. pipiens* at 10°C, below the 14°C lower limit of previous work[34]. Surprisingly, dissemination at 10°C (24– 27%) for both virus strains exceeded that at 15°C (∼6%) and prior reported peaks (∼10%)[2], suggesting complex interactions between viral replication and host physiology at cooler temperatures. One possible mechanism is temperature-dependent suppression of RNA interference (RNAi), a central antiviral pathway in mosquitoes, consistent with observations in other arboviruses such as chikungunya and yellow fever viruses[35]. Semi-mechanistic modeling incorporating a temperature-dependent tradeoff between viral replication and mosquito immunity produced a bimodal competence thermal performance curve consistent with our empirical data, supporting the plausibility of this mechanism. While this non-unimodality did not markedly alter overall thermal predictions of WNV transmission in our system, because other mosquito traits limit transmission at these low temperatures, it raises the possibility that bimodal transmission patterns may emerge in other mosquito–pathogen systems and should be considered in future ecologically informed modeling efforts. Moreover, under more complex variable temperature scenarios, elevated vector competence at low temperature could combine with mosquito overwintering or life history variation to allow transmission—a possibility that could merit further modeling or empirical work.

Here, we highlight the importance of incorporating transmission-relevant data, specifically life-history traits of WNV-infected mosquitoes, into models of arbovirus risk. We detected differences in the relative R₀ between unexposed and WNV-infected mosquitoes, as well as strain-specific responses at higher temperatures. While thermal limits and optimum of relative R₀ did not differ significantly between models using traits only from unexposed versus transmission-relevant WNV-infected mosquito traits, we observed substantial changes in the overall level of transmission risk. These results demonstrate that accounting for the effects of WNV infection on mosquito life-history traits provides a more accurate prediction of WNV risk and its temperature-dependent dynamics. This framework also sets the stage for future ecological investigations of other arbovirus–mosquito systems under natural field conditions.

## Acknowledgements

We thank the NYS Arbovirology Laboratory insectary staff for support and assistance. We thank the Wadsworth Center Media and Tissue Culture Facility for providing cells and media.

## Contributions

RLF and ATC conceived and designed the study. RLF and EMB collected and formatted the data utilized in the manuscript. MCL developed and modified models used in this work. RLF and MCL conducted the results analysis. RLF, MCL, EAM, and ATC prepared the manuscript. EAM and ATC supervised the work. RLF, MCL, EMB, EAM, and ATC provided constructive comments to improve the manuscript and participated in revising the manuscript.

