## Appendix for "Mosquito infection-mediated trait variation alters temperature-dependent transmission of West Nile virus"

#### Temperature-dependent relative $\mathcal{R}_0$

##### Derivation of a relative $\mathcal{R}_0$ model based on combined traits of unexposed and infected mosquitoes

The Ross-MacDonald model for mosquito transmitted diseases yields the following estimate of the basic reproductive number $\mathcal{R}_0$:

$$\mathcal{R}_0 = \sqrt{\frac{a^2(T)b(T)c(T)e^{-\frac{\mu(T)}{\mathrm{PDR}(T)}}M(T)}{Nr\mu(T)}} \tag{1}$$

where $a(T)$ is the biting rate (in units of bites per mosquito per day), $b(T)$ the probability that a mosquito becomes infected after biting an infected host, $c(T)$ the probability that a host becomes infected after being bitten by an infected mosquito, $\mu(T)$ the death rate of mosquitoes, $\mathrm{PDR}(T)$ the pathogen development rate (the inverse of the extrinsic incubation period), $M(T)$ the number of mosquitoes, $N$ the number of hosts and $r$ the host recovery rate.

Shocket *et al* use the following approximation for mosquito abundance based on various temperature-dependent life history traits [1]:

$$\begin{aligned} M(T) &\approx \frac{\mathrm{EFD}(T)\mathrm{EV}(T)p_{LA}(T)\mathrm{MDR}(T)}{\mu^2(T)} \\ &= \frac{a_G(T)\mathrm{EFGC}(T)\mathrm{EV}(T)p_{LA}(T)\mathrm{MDR}(T)}{\mu^2(T)} \end{aligned}$$

where $\mathrm{EFD}(T)$ is the number of eggs per female per day, $\mathrm{EV}(T)$ the egg viability, $p_{LA}(T)$ the probability that a larva survives to adulthood, $\mathrm{MDR}(T)$ the mosquito development rate and $\mu^2(T)$ the death rate. The number of eggs laid per female can be expressed as the number of eggs per female per gonotrophic cycle $\mathrm{EFGC}(T)$ times the number of gonotrophic cycles per day $a_G(T)$ (i.e. inverse of the gonotrophic cycle length). The notation $a_G$ is used because the biting rate corresponds to the inverse of the gonotrophic cycle length under the assumption that a mosquito will feed exactly once per gonotrophic cycle, which has been used in previous models of WNV transmission [1]. However, this assumption may not be entirely accurate, as mosquitoes can occassionally feed multiple times depending on experimental conditions. Because of this, here we distinguish between $a_G(T)$ and the true biting rate $a(T)$, which can be estimated directly through mosquito bloodfeeding experiments.

In this section, show how this model can be modified to account for mosquito traits that vary with infection status. First, it is helpful to factorize the terms inside the square root to better understand how the expression for $\mathcal{R}_0$ can be derived (following a similar approach as in [2]).

$$\mathcal{R}_0 = \sqrt{\left[M(T)a(T)\left(\frac{1}{N}\right)\left(\frac{1}{r}\right)b(T)\right]e^{-\mu(T)\left(\frac{1}{PDR(T)}\right)}\left[a(T)\left(\frac{1}{\mu(T)}\right)c(T)\right]} \tag{2}$$

The quantity inside the square root in Equation 1 consists of the number of secondary cases arising from a single infected host. The first term in brackets corresponds to the total number of mosquitoes that become infected from a single infected host. This can be broken down as follows:

- $M(T)a(T)$ is the total number of mosquito bites per day.
- Assuming all hosts are equally likely to be bitten, the probability of biting the infected host is $\frac{1}{N}$. The length of time that the host is infectious is the inverse of the recovery rate $\frac{1}{r}$ (where $r$ is in units of day$^{-1}$). So $M(T)a(T)\left(\frac{1}{N}\right)\left(\frac{1}{r}\right)$ is the total number of mosquitoes that bite the infected host during the period where it is infective.
- This quantity is then multiplied by $b(T)$, the probability that a mosquito becomes infected after biting an infected host. This yields the total number of infected *mosquitoes* arising from an initial single infected host.

An infected mosquito is not immediately infectious. The virus must first develop and spread within the mosquito, eventually reaching the salivary glands after an *extrinsic incubation period* $t_{\text{EIP}}(T)$. Due to their short lifespan, a significant fraction of mosquitoes can die before becoming infectious. This model accounts for this with the factor $e^{-\mu(T)t_{\text{EIP}}(T)} = e^{-\frac{\mu(T)}{\text{PDR}(T)}}$ which represents the fraction of surviving mosquitoes after the extrinsic incubation period. The inverse of the extrinsic incubation period is the pathogen development rate $\text{PDR}(T)$, which has a unimodal relationship with temperature and can be modeled with standard equations for thermal performance curves. The product of the first term in brackets and the factor $e^{-\frac{\mu(T)}{\text{PDR}(T)}}$ in Equation 2 thus corresponds to the total number of infected mosquitoes that survive past the extrinsic incubation period (and are infectious).

The third term in parenthesis corresponds to the number of infected hosts that arise from an infectious mosquito. It can be broken down as follows:

- $a(T)\left(\frac{1}{\mu(T)}\right)$ corresponds to the total number of bites per infectious mosquito during the period that the mosquito remains infectious. Mosquitoes are assumed to not recover from West Nile Virus. However, they will eventually die, at which point they cannot transmit WNV anymore. The death rate is thus used here in lieu of a recovery rate.

- If we assume all bites are on susceptible hosts (a reasonable assumption if $N$ is large), a fraction $c(T)$ of these bites will result on hosts becoming infected with WNV.

The total number of secondary infectious hosts arising from an initial single infected host is thus given by the product of the terms underneath the square root. The square root is sometimes included to account for the number of secondary hosts per transmission event (as two mosquito bites are necessary: one to transmit the virus to a mosquito and one to transmit it to a new host). Multiplying through these terms leads to the usual Ross-MacDonald model (Equation 1).

##### Incorporating life history traits that vary in infected mosquitoes

If we allow for infectious mosquitoes to have a different biting rate and death rate when infected by West Nile virus (denoted by $a_I$ and $\mu_I$) , the derivation can be adjusted as follows:

$$\mathcal{R}_0 = \sqrt{\left[M(T)a(T)\left(\frac{1}{N}\right)\left(\frac{1}{r}\right)b(T)\right]e^{-\mu_I(T)\left(\frac{1}{PDR(T)}\right)}\left[a_I(T)\left(\frac{1}{\mu_I(T)}\right)c(T)\right]}$$
$$= \sqrt{\frac{a(T)a_I(T)b(T)c(T)e^{-\frac{\mu_I(T)}{PDR(T)}}M(T)}{Nr\mu_I(T)}} \tag{3}$$

The first term in brackets in Equation 3 remains unchanged, as it corresponds to bites from mosquitoes that are not infected with WNV. The infected death rate and biting rate are used in the rest of the terms, as they correspond to survival of infected mosquitoes through the extrinsic incubation period and the number of bites of infectious mosquitoes. This updated model allows us to include the effects of infection in the relevant life history traits for transmission, and reduces to the usual model if these effects do not exist, as in this case we have $a(T) = a_I(T)$ and $\mu_I(T) = \mu(T)$.

In practice, it is difficult to estimate the number of hosts $N$ and the host recovery rate $r$. However, these traits are assumed to be temperature-independent and thus are not expected to modify the $\mathcal{R}_0$ curve. Because of this, it is common to remove these quantities from the expressions above, yielding a relative $\mathcal{R}_0$ that is proportional to transmission but that has an arbitrary scale that does not carry the same interpretation in terms of secondary cases [1]. We follow the same approach here to develop a relative $\mathcal{R}_0$ with either exposed or infected traits, simply removing $N$ and $r$ from the expression for $\mathcal{R}_0$ (see Methods).

### Thermal performance curves

In this section, we describe the methods used to estimate thermal performance curves for the life history traits of *Culex pipiens*, both in the absence of WNV, and when exposed/infected with two strains of West Nile Virus (2003.2 and 2017.1). For many traits, we observed that the shape of the thermal performance curve changed in infected mosquitoes. To facilitate comparisons of the TPCs in these cases, we used flexTPC [3], a flexible model for thermal performance curves that allows direct comparisons across conditions with TPCs of different shapes:

$$r(T) = r_{\max}\left[\left(\frac{T - T_{\min}}{\alpha}\right)^{\alpha}\left(\frac{T_{\max} - T}{1-\alpha}\right)^{1-\alpha}\left(\frac{1}{T_{\max} - T_{\min}}\right)\right]^{\frac{\alpha(1-\alpha)}{\beta^2}}$$

The optimal temperature in the flexTPC model is given by

$$T_{\text{opt}} = \alpha T_{\max} + (1-\alpha)T_{\min}$$

#### Vector competence

One of the most striking features of the data presented here is finding vector competence thermal performance curves that are clearly not unimodal. While this could have multiple explanations, we hypothesize that these curves may arise due to an interplay between the thermal dependence of viral replication and mosquito immunity. The unusual shape of the TPCs found in this work made it necessary to develop a mathematical model that can describe them. We describe the development of this novel mathematical model, based on the replication/immunity tradeoff hypothesis, in this section.

As a simple approximation to the interplay of viral replication rates and mosquito immunity, we model the initial stages of replication of West Nile virus in an exposed mosquito with a simple population dynamics model. We represent the amount of virus $v$ in a mosquito with the differential equation

$$\frac{dv}{dt} = r_v(T)v\left(1 - \frac{v}{K}\right) - i_m(T)v$$

where $r_v(T)$ is the viral replication rate and $i_m(T)$ the rate at which viral particles are cleared by mosquito immunity (both of which are functions of temperature $T$) and $K$ is the carrying capacity of virus (i.e. the maximum number of viral particles that can be supported by a mosquito where immunity is completely nonfunctional). At steady state, this equation implies that the amount of virus will be

$$v^*(T) = K\left(1 - \frac{i_m(T)}{r_v(T)}\right) \tag{4}$$

We will call $v^*$ the *viral load*, which depends on the balance of the viral growth rate and mosquito immunity in a temperature-sensitive manner. For simplicity, as no direct measurements are made of the number of viral particles, we will choose the units so that the carrying capacity $K = 1$. This is equivalent to making the units of virus a proportion relative to the number of viral particles that are produced when the mosquito immunity is completely nonfunctional (i.e. when $i_m(T) = 0$).

Next, we assume that the probability that the infection progresses (and is detectable in the mosquito carcass) increases monotonically with the viral load. Likewise, the probability that an infection progresses to a disseminated infection (when the virus is detected in the legs) is also assumed to increase monotonically with the viral load. For simplicity, we represent this with power functions

$$b(T) \propto \left[\frac{v^*(T)}{K}\right]^{n_b} = [v^*(T)]^{n_b}$$
$$c(T) \propto \left[\frac{v^*(T)}{K}\right]^{n_c} = [v^*(T)]^{n_c}$$

(where the equalities are true because of setting $K = 1$). Moreover, note that it must be true that when $v^* = 0$ (i.e. when there is no virus), there are no infected or disseminated mosquitoes. We assume that when $v^* = 1$ (i.e. when there is nonfunctioning immunity and the viral load is maximized), all mosquitoes develop a disseminated infection ($b(T) = c(T) = 1$). Under these assumptions, we can replace the proportionality signs with equalities

$$b(T) = [v^*(T)]^{n_b}$$
$$c(T) = [v^*(T)]^{n_c}$$

Also note that, by definition, the probability of a disseminated infection $bc(T)$ will be

$$bc(T) = b(T)c(T) = [v^*(T)]^{n_b+n_c} = [v^*(T)]^{n_{bc}}$$

where $n_{bc} = n_b + n_c$ (also note that, since $v^* \in [0, 1]$, $b(T), c(T), bc(T) \in [0, 1]$).

For increased interpretability, we reparameterize the model by replacing the exponents $n_b$ and $n_{bc}$ with the viral load that leads to 50% of the mosquitoes being infected ($v^*_{50,b}$) or disseminated ($v^*_{50,bc}$), respectively. This can be done by solving the equations $b(T) = \frac{1}{2}$ and $bc(T) = \frac{1}{2}$. For example, for $b(T)$ we have.

$$\frac{1}{2} = \left[v^*_{50,b}\right]^{n_b}$$
$$n_b \ln v^*_{50} = \ln \frac{1}{2}$$
$$n_b = -\frac{\ln 2}{\ln v^*_{50,b}}$$

Repeating this procedure replacing $b$ with $bc$ leads to $n_{bc} = -\frac{\ln 2}{\ln v^*_{50,bc}}$ where $v^*_{50,bc}$. Replacing $n_b$ and $n_{bc}$ in the expressions for $b(T)$ and $bc(T)$ leads to

$$\begin{aligned} b(T) &= v^*(T)^{-\frac{\ln 2}{\ln v^*_{50,b}}} \\ bc(T) &= v^*(T)^{-\frac{\ln 2}{\ln v^*_{50,bc}}} \end{aligned} \tag{5}$$

Disseminated infections are a subset of infections. Because of this, we assume that the viral load needed to have 50% dissemination is larger than or equal that the one needed to have 50% infection. In other words, the inequality $v^*_{50,bc} \geq v^*_{50,b}$ is a constraint that must be satisfied.

We model WNV clearance rate due to mosquito immunity with a Hill equation

$$i_m(T) = i_0 + (i_{\max} - i_0)\frac{T^n}{T_{50}^n + T^n}$$

which assumes that the immunity clearance rate decreases to a minimum value $i_0$ at 0°C but plateaus at high temperatures at a value $i_{\max}$. Here, $T_{50}$ is the temperature at which the viral clearing rate because of immunity is 50% of the maximum and $n$ is the Hill exponent which determines the steepness of the immunity thermal response.

The initial viral replication rate inside the mosquito was modeled as a unimodal function with a flexTPC functional form:

$$r_v(T) = r_{v,\max}\left[\left(\frac{T - T_{\min}}{\alpha}\right)^{\alpha}\left(\frac{T_{\max} - T}{1-\alpha}\right)^{1-\alpha}\left(\frac{1}{T_{\max} - T_{\min}}\right)\right]^{\frac{\alpha(1-\alpha)}{\beta^2}}$$

where $r_{v,\max}$ is the maximum viral replication rate, $T_{\min}$ the minimum temperature and $T_{\max}$ the maximum temperature, $\alpha \in (0,1)$ the location of the thermal optimum relative to the maximum and minimum and $\beta$ determines the thermal breadth.

We make some further simplifying assumptions to reduce the number of parameters. First, note that $v^*(T)$ depends only on the ratio $\frac{i_m(T)}{r_v(T)}$. Because of this, we can reduce one parameter by setting $i_{\max} = 1$, making $i_m(T)$ a proportion relative to the maximum immunity clearance rate. This still allows us to find the needed ratio if we choose the units of the viral replication rate $r_v(T)$ to be relative to this maximum immunity clearance rate. Lastly, to reduce the number of parameters we set $\alpha = 0.8$ and $\beta = 0.2$, which assumes that the initial viral replication rate TPCs are left-skewed curves with a typical thermal breadth.

##### Statistical model

The number of infected mosquitoes for each strain was modeled with a binomial distribution. Four thermal performance curves (proportion infected and disseminated for two strains) were inferred simultaneously, with common parameters for mosquito immunity (shared for all four curves), per-strain parameters for the viral replication rate (shared for )

$$
\begin{aligned}
n_{\text{inf},s,T}|T, b(T) &\sim \text{Binomial}(b_s(T), N_T) \\
n_{\text{dis},s,T}|T, bc(T) &\sim \text{Binomial}(bc_s(T), N_T) \\
b_s(T)|v_{50,b,s}, v^*(T) &= v^*(T)^{-\frac{\ln 2}{v_{50,b,s}}} \\
bc_s(T)|v_{50,bc,s}, v^*(T) &= v^*(T)^{-\frac{\ln 2}{v_{50,bc,s}}} \\
v^*(T)|r_v(T), i_m(T) &= 1 - \frac{i_m(T)}{r_v(T)} \\
i_m(T)|T, T_{50}, n, i_0 &= i_0 + (1 - i_0)\frac{T^n}{T_{50}^n + T^n} \\
r_{v,s}(T)|T_{\min,s}, T_{\max,s}, r_{\max,s}, \alpha, \beta &= r_{\max,s}\left[\left(\frac{T - T_{\min,s}}{\alpha}\right)\left(\frac{T_{\max,s} - T}{1 - \alpha}\right)\left(\frac{1}{T_{\max,s} - T_{\min,s}}\right)\right]^{\frac{\alpha(1-\alpha)}{\beta^2}}
\end{aligned}
$$

##### Prior distributions

The priors for the mosquito immunity parameters were:

$$
\begin{aligned}
T_{50} &\sim \text{Normal}(\mu = 20°\text{C}, \sigma = 10°\text{C}) \\
\log n &\sim \text{Normal}(\mu = 0, \sigma = 0.5) \\
i_0 &\sim \text{Uniform}(0, 1)
\end{aligned}
$$

These are weak priors that assume that the temperature of half-maximum immunity is approximately 95% likely *a priori* to be in the interval $[0°\text{C}, 40°\text{C}]$, that the Hill exponent is 95% likely *a priori* to be in the interval $[0.1, 10]$ and that the rate of WNV clearance due to immunity at 0°C is equally likely to be any value in-between 0% and 100% of the maximum clearance rate at high temperatures.

The following priors were used for the initial viral replication rate $r_{v,s}$:

$$
\begin{aligned}
T_{\min,s} &\sim \text{Normal}(\mu = 5°\text{C}, \sigma = 2.5°\text{C}) \\
T_{\max,s} &\sim \text{Normal}(\mu = 35°\text{C}, \sigma = 5°\text{C}) \\
r_{\max,s} &\sim \text{Exponential}(\mu = 1)
\end{aligned}
$$

A separate TPC was fit for each strain, with identical priors. As described earlier, to reduce the number of parameters we set $\alpha = 0.8$ and $\beta = 0.2$.

The following priors were used for the viral loads leading to 50% infection and dissemination:

$$
\begin{aligned}
v_{50,bc,s}|v_{50,b,s} &\sim \text{TruncatedBeta}(a = 9, b = 21, a = v_{50,b,s}, b = 1) \\
v_{50,b,s} &\sim \text{Beta}(a = 6, b = 24)
\end{aligned}
$$

As no direct measurements of the viral load were made, these priors were chosen to be informative in order to make the model identifiable. One assumption made here is that the viral loads that would be reached with no immunity are much higher than those in its presence. For example, the prior for $v_{50,b,s}$ assumes that the viral load that results in 50% mosquito infection is around 20% (95% prior CI: $[8\%, 37.8\%]$) of the maximum possible viral load (attained when immunity is completely non-functional).The prior for $v_{50,bc,s}$ is a truncated Beta prior that enforces the requirement that $v_{50,bc,s} \geq v_{50,b,s}$.

**Posterior summaries**

| Trait | Strain | $N_{\text{TPC}}$ | Parameter | Mean | SD | 2.5% | 97.5% | $\hat{R}$ | $n_{\text{eff}}$ |
|---|---|---|---|---|---|---|---|---|---|
| $b$ | 2017.1 | 1 | $v_{50,b}$ | 0.124 | 0.038 | 0.061 | 0.210 | 1.001 | 75000 |
| $b$ | 2003.2 | 1 | $v_{50,b}$ | 0.222 | 0.054 | 0.129 | 0.338 | 1.001 | 52000 |
| $bc$ | 2017.1 | 1 | $v_{50,bc}$ | 0.398 | 0.054 | 0.295 | 0.504 | 1.001 | 58000 |
| $bc$ | 2003.2 | 1 | $v_{50,bc}$ | 0.416 | 0.056 | 0.309 | 0.528 | 1.001 | 60000 |
| $r_v$ | 2017.1 | 2 | $T_{\max}$ | 40.8 | 2.4 | 37.1 | 46.2 | 1.001 | 50000 |
| $r_v$ | 2017.1 | 2 | $T_{\min}$ | 1.0 | 1.4 | -2.0 | 3.6 | 1.001 | 11000 |
| $r_v$ | 2017.1 | 2 | $T_{\text{opt}}$ | 32.9 | 1.8 | 30.0 | 37.0 | 1.001 | 58000 |
| $r_v$ | 2017.1 | 2 | $r_{\max}$ | 3.52 | 0.58 | 2.59 | 4.87 | 1.001 | 35000 |
| $r_v$ | 2003.2 | 2 | $T_{\max}$ | 35.0 | 0.4 | 34.4 | 36.0 | 1.001 | 100000 |
| $r_v$ | 2003.2 | 2 | $T_{\min}$ | 1.8 | 1.3 | -0.8 | 4.2 | 1.001 | 12000 |
| $r_v$ | 2003.2 | 2 | $T_{\text{opt}}$ | 28.4 | 0.3 | 27.8 | 29.1 | 1.001 | 41000 |
| $r_v$ | 2003.2 | 2 | $r_{\max}$ | 2.51 | 0.32 | 1.96 | 3.21 | 1.001 | 47000 |
| $i_m$ | all | 4 | $T_{50}$ | 14.2 | 0.9 | 12.4 | 15.8 | 1.001 | 12000 |
| $i_m$ | all | 4 | $n$ | 7.357 | 3.029 | 5.082 | 14.387 | 1.001 | 25000 |
| $i_m$ | all | 4 | $i_{\min}$ | 0.056 | 0.052 | 0.001 | 0.190 | 1.001 | 31000 |

#### Biting rate

Biting rate $a(T)$ was modeled with a flexTPC curve

$$a(T) = a_{\max} \left[ \left( \frac{T - T_{\min}}{\alpha} \right)^{\alpha} \left( \frac{T_{\max} - T}{1 - \alpha} \right)^{1-\alpha} \left( \frac{1}{T_{\max} - T_{\min}} \right) \right]^{\frac{\alpha(1-\alpha)}{\beta^2}}$$

where $a_{\max}$ is the maximum biting rate, $T_{\min}$ and $T_{\max}$ the minimum and maximum temperatures, respectively, $\alpha \in (0, 1)$ a parameter that determines the position of the thermal optimum relative to the minimum and maximum (where $T_{\text{opt}} = \alpha T_{\max} + (1-\alpha) T_{\min}$) and $\beta$ a parameter that represents the approximate ratio of the thermal breadth at 88% maximum performance and the thermal tolerance range $T_{\max} - T_{\min}$. A separate curve was fit to mosquitoes unexposed to WNV, mosquitoes exposed to the 2003.2 or 2017.1 strain (but uninfected), and mosquitoes infected with the 2003.2 or 2017.1 strain.

##### Statistical model

##### Likelihood

The biting rate of a mosquito is defined as the number of bites (assumed to be equal to the number of bloodfeeding events) per mosquito per unit time. In the experiments, a bloodmeal was offered to the mosquitoes weekly. Thus, the proportion of accepted bloodmeals (i.e. number of feeding events divided by the number of offered bloodmeals) corresponds to the weekly biting rate.

The number of bloodfeeding events $n_i$ of mosquito $i$ under experimental treatment $c$ (which can be one of unexposed, exposed or infected with a WNV strain) and temperature $T_i$ was modeled with a binomial distribution

$$n_{ci}|T_i, \mathcal{P}_c \sim \text{Binomial}(p = a_c(T_i, \mathcal{P}_c), N_{ci})$$

where $\mathcal{P}_c = \{T_{\min}^{(c)}, T_{\max}^{(c)}, a_{\max}^{(c)}, \alpha^{(c)}, \beta^{(c)}\}$ are the parameters of a flexTPC equation for experimental group $c$.

As the biting rate is assumed to be the same for all mosquitoes of the same experimental group that were kept at the same temperature, this model is equivalent to modeling the total number of accepted bloodmeals of mosquitoes of group $c$ at temperature $T$ $n_{cT} = \sum_i n_{ci} 1_{\{T_i = T\}}$ with a binomial distribution

$$n_{cT}|\mathcal{P}_c \sim \text{Binomial}(p = a_c(T, \mathcal{P}_c), N_{cT})$$

where $N_{cT} = \sum_i N_{ci} 1_{\{T_i = T\}}$ is the total number of offered bloodmeals to mosquitoes of the same experimental group and temperature treatment. The latter form of the model was used for inference for computational efficiency.

##### Prior distributions for unexposed mosquitoes

In previous work, biting rate has been estimated from the inverse of the gonotrophic cycle length, which yielded a left-skewed TPC. In contrast, here the biting rate is estimated directly from bloodfeeding experiments. The shape of the curve looks qualitatively different, with our data being consistent with a more symmetric curve. Because of this, we did not use informative priors based on the previously estimated curves for this trait.

Instead, for the unexposed mosquitoes, we use priors that are weakly informative and only constrain the parameters to biologically reasonable ranges. The following prior distributions were used for the flexTPC parameters for the biting rate of unexposed mosquitoes:

$$\begin{aligned}
T_{\min} &\sim \text{Normal}(\mu = 7.5\ {}^\circ\text{C}, \sigma = 3.75\ {}^\circ\text{C}) \\
T_{\max} &\sim \text{Normal}(\mu = 35\ {}^\circ\text{C}, \sigma = 3\ {}^\circ\text{C}) \\
a_{\max} &\sim \text{Uniform}(0, 1) \\
\alpha &\sim \text{Uniform}(0, 1) \\
\beta &\sim \text{Gamma}(\mu = 0.35, \sigma = 0.2)
\end{aligned}$$

The prior for $T_{\text{min}}$ was chosen to have approximately 95% prior probability of the minimum temperature being in the interval $[0\ ^\circ\text{C}, 15\ ^\circ\text{C}]$. The prior for $T_{\text{max}}$ was chosen so an approximate prior 95% CI is $[29\ ^\circ\text{C}, 41\ ^\circ\text{C}]$, The prior for $a_{\text{max}}$ assumes that any feeding proportion is equally likely *a priori*. Likewise, the prior for $\alpha$ is a uniform prior assumes that the optimum temperature is equally likely to be at any point in-between $T_{\text{min}}$ and $T_{\text{max}}$. The prior for $\beta$ is weakly informative, preferring a thermal breadth similar to those in common TPC models like the Briere and quadratic models, but allowing for other values if necessary to describe the data.

**Posterior summaries for unexposed mosquitoes**

| Parameter | Mean | SD | 2.5% | 97.5% | $\hat{R}$ | $n_{\text{eff}}$ |
|---|---|---|---|---|---|---|
| $T_{\text{min}}$ | 9.4 | 3.1 | 2.5 | 14.4 | 1.001 | 75000 |
| $T_{\text{opt}}$ | 22.5 | 1.0 | 20.5 | 24.4 | 1.001 | 58000 |
| $T_{\text{max}}$ | 34.7 | 2.4 | 30.8 | 40.0 | 1.001 | 31000 |
| $a_{\text{max}}$ | 0.307 | 0.046 | 0.223 | 0.401 | 1.001 | 38000 |
| $\alpha$ | 0.513 | 0.099 | 0.309 | 0.699 | 1.001 | 67000 |
| $\beta$ | 0.210 | 0.062 | 0.124 | 0.366 | 1.001 | 36000 |

**Prior distributions for exposed and infected mosquitoes**

We initially attempted to use the same prior distributions for the exposed and infected groups. However, some group/temperature combinations had a very low number of mosquitoes, which led to the TPCs being poorly identified. Because of this, we decided to use informative priors based on the curves fit to the unexposed mosquitoes. This approach is conservative, as the priors will pull the estimates towards those of the unexposed mosquitoes.

$$\begin{aligned}
T_{\text{min}} &\sim \text{Normal}(\mu = 9.370\ ^\circ\text{C}, \sigma = 3.5\ ^\circ\text{C}) \\
T_{\text{max}} &\sim \text{Normal}(\mu = 34.716\ ^\circ\text{C}, \sigma = 3\ ^\circ\text{C}) \\
a_{\text{max}} &\sim \text{Uniform}(0, 1) \\
\alpha &\sim \text{Beta}(\mu = 0.513, \sigma = 0.15) \\
\beta &\sim \text{Gamma}(\mu = 0.21, \sigma = 0.062)
\end{aligned}$$

The priors for $T_{\text{min}}$, $T_{\text{max}}$ and $\alpha$ are informative, and based on the estimates for unexposed mosquitoes but with increased variance to account for the fact that these parameters may be different in these experimental groups. The prior for $\beta$ is also informative, but with the same standard deviation as the posterior estimate for the unexposed mosquitoes (i.e. without inflating the variance). This variance was not inflated as initial attempts to fit the model with an inflated standard deviation yielded some curves that were unrealistically narrow, with a very high peak in-between measurements and rapidly decreasing biting rate from the peak.

**Posterior summaries for exposed and infected mosquitoes**

**2003.2 exposed uninfected**

| Parameter | Mean | SD | 2.5% | 97.5% | $\hat{R}$ | $n_{\text{eff}}$ |
|---|---|---|---|---|---|---|
| $T_{\text{min}}$ | 9.7 | 2.5 | 4.2 | 13.8 | 1.001 | 75000 |
| $T_{\text{opt}}$ | 26.1 | 2.2 | 22.2 | 31.1 | 1.001 | 65000 |
| $T_{\text{max}}$ | 35.6 | 2.5 | 31.3 | 40.9 | 1.001 | 50000 |
| $a_{\text{max}}$ | 0.196 | 0.073 | 0.079 | 0.362 | 1.001 | 75000 |
| $\alpha$ | 0.635 | 0.103 | 0.429 | 0.830 | 1.001 | 54000 |
| $\beta$ | 0.206 | 0.048 | 0.129 | 0.316 | 1.001 | 75000 |

**2003.2 infected**

| Parameter | Mean | SD | 2.5% | 97.5% | $\hat{R}$ | $n_{\text{eff}}$ |
|---|---|---|---|---|---|---|
| $T_{\text{min}}$ | 10.8 | 3.1 | 4.3 | 16.6 | 1.001 | 46000 |
| $T_{\text{opt}}$ | 23.9 | 2.0 | 19.9 | 28.0 | 1.001 | 67000 |
| $T_{\text{max}}$ | 34.1 | 2.8 | 29.1 | 39.7 | 1.001 | 75000 |
| $a_{\text{max}}$ | 0.103 | 0.045 | 0.045 | 0.196 | 1.001 | 75000 |
| $\alpha$ | 0.561 | 0.110 | 0.341 | 0.775 | 1.001 | 70000 |
| $\beta$ | 0.191 | 0.055 | 0.100 | 0.316 | 1.001 | 75000 |

**2017.1 exposed uninfected**

| Parameter | Mean | SD | 2.5% | 97.5% | $\hat{R}$ | $n_{\text{eff}}$ |
|---|---|---|---|---|---|---|
| $T_{\text{min}}$ | 9.4 | 2.7 | 3.4 | 13.9 | 1.001 | 72000 |
| $T_{\text{opt}}$ | 22.3 | 3.1 | 16.9 | 29.1 | 1.001 | 75000 |
| $T_{\text{max}}$ | 34.2 | 3.0 | 28.5 | 40.2 | 1.001 | 75000 |
| $a_{\text{max}}$ | 0.090 | 0.058 | 0.019 | 0.240 | 1.001 | 42000 |
| $\alpha$ | 0.518 | 0.126 | 0.276 | 0.773 | 1.001 | 49000 |
| $\beta$ | 0.201 | 0.057 | 0.107 | 0.328 | 1.001 | 75000 |

**2017.1 infected**

| Parameter | Mean | SD | 2.5% | 97.5% | $\hat{R}$ | $n_{\text{eff}}$ |
|---|---|---|---|---|---|---|
| $T_{\text{min}}$ | 8.3 | 2.7 | 2.5 | 13.0 | 1.001 | 41000 |
| $T_{\text{opt}}$ | 25.6 | 2.4 | 20.8 | 30.5 | 1.001 | 75000 |
| $T_{\text{max}}$ | 35.7 | 2.5 | 31.4 | 41.0 | 1.001 | 26000 |
| $a_{\text{max}}$ | 0.081 | 0.024 | 0.042 | 0.136 | 1.001 | 75000 |
| $\alpha$ | 0.634 | 0.115 | 0.398 | 0.849 | 1.001 | 28000 |
| $\beta$ | 0.254 | 0.056 | 0.162 | 0.381 | 1.001 | 75000 |

#### Adult lifespan

In Shocket *et al*, lifespan was modeled with a piece-wise linear model. Here, we model lifespan as a unimodal TPC instead. This is arguably more realistic as there will eventually be a temperature that is too low for the mosquitoes to survive. Adult lifespan $\mathrm{lf}(T) = \frac{1}{\mu(T)}$ was modeled with a flexTPC curve

$$\mathrm{lf}(T) = \mathrm{lf}_{\max}\left[\left(\frac{T - T_{\min}}{\alpha}\right)^{\alpha}\left(\frac{T_{\max} - T}{1-\alpha}\right)^{1-\alpha}\left(\frac{1}{T_{\max} - T_{\min}}\right)\right]^{\frac{\alpha(1-\alpha)}{\beta^2}}$$

where $\mathrm{lf}_{\max}$ is the maximum lifespan, $T_{\min}$ and $T_{\max}$ the minimum and maximum temperatures, respectively, $\alpha \in (0,1)$ a parameter that determines the position of the thermal optimum relative to the minimum and maximum (where $T_{\mathrm{opt}} = \alpha T_{\max} + (1-\alpha)T_{\min}$) and $\beta$ a parameter that represents the approximate ratio of the thermal breadth at 88% maximum performance and the thermal tolerance range $T_{\max} - T_{\min}$. A separate curve was fit to mosquitoes unexposed to WNV, mosquitoes exposed to the 2003.2 or 2017.1 strain (but uninfected), and mosquitoes infected with the 2003.2 or 2017.1 strain.

##### Statistical model

Lifespan was modeled with a negative binomial distribution.

$$y_i | T_i, \mathcal{P}, r \sim \mathrm{NegativeBinomial}(\mu = \mathrm{lf}(T; \mathcal{P}), r = r)$$

##### Prior distributions

$$\begin{aligned} T_{\min} &\sim \mathrm{Normal}(\mu = 5\ ^\circ\mathrm{C}, \sigma = 2.5\ ^\circ\mathrm{C}) \\ T_{\max} &\sim \mathrm{Normal}(\mu = 34.9\ ^\circ\mathrm{C}, \sigma = 2.5\ ^\circ\mathrm{C}) \\ \mathrm{lf}_{\max} &\sim \mathrm{Uniform}(0, 150) \\ \alpha &\sim \mathrm{Uniform}(0, 1) \\ \beta &\sim \mathrm{Gamma}(\mu = 0.35, \sigma = 0.2) \\ r &\sim \mathrm{Uniform}(0, 50) \end{aligned}$$

The prior we chose for the minimum temperature is a weakly informative, but biologically conservative. It assumes the minimum temperature is a priori approximately 95% likely to be in the interval $[0\ ^\circ\mathrm{C}, 10\ ^\circ\mathrm{C}]$. The prior for the maximum temperature is a weakly informative prior based on a previous estimate for the maximum lifespan of *Cx pipiens* in Shocket *et al.* It has an inflated variance to allow for the possible differences in the mosquito populations in this work compared to previous studies, with an approximate prior 95% CI of $[29.9\ ^\circ\mathrm{C}, 39.9\ ^\circ\mathrm{C}]$. The prior for the maximum lifespan assumes all values between 0 and 150 days are equally likely *a priori*, where the upper limit was chosen based on previous data for *Cx pipiens*. The prior for $\alpha$ is a uniform prior that assumes that the optimum temperature is equally likely to be at any point in-between $T_{\min}$ and $T_{\max}$. The prior for $\beta$ is weakly informative, preferring a thermal breadth similar to those in common TPC models like the Briere and quadratic models, but allowing for other values if necessary to describe the data..

##### Posterior summaries

##### Unexposed mosquitoes

| Parameter | Mean | SD | 2.5% | 97.5% | $\hat{R}$ | $n_{\text{eff}}$ |
|---|---|---|---|---|---|---|
| $T_{\min}$ | 5.4 | 2.2 | 0.7 | 9.3 | 1.001 | 52000 |
| $T_{\text{opt}}$ | 20.1 | 1.2 | 17.5 | 22.4 | 1.002 | 38000 |
| $T_{\max}$ | 36.0 | 1.6 | 33.5 | 39.7 | 1.001 | 48000 |
| $\text{lf}_{\max}$ | 21.5 | 1.6 | 18.6 | 24.8 | 1.001 | 180000 |
| $\alpha$ | 0.480 | 0.077 | 0.316 | 0.620 | 1.001 | 43000 |
| $\beta$ | 0.397 | 0.075 | 0.282 | 0.572 | 1.001 | 32000 |
| $r$ | 2.86 | 0.36 | 2.22 | 3.61 | 1.001 | 250000 |

##### 2003.2 exposed uninfected

| Parameter | Mean | SD | 2.5% | 97.5% | $\hat{R}$ | $n_{\text{eff}}$ |
|---|---|---|---|---|---|---|
| $T_{\min}$ | 4.1 | 2.3 | -0.5 | 8.4 | 1.001 | 210000 |
| $T_{\text{opt}}$ | 7.8 | 2.9 | 2.2 | 13.5 | 1.001 | 96000 |
| $T_{\max}$ | 35.7 | 1.7 | 33.3 | 39.5 | 1.001 | 180000 |
| $\text{lf}_{\max}$ | 16.1 | 1.6 | 13.5 | 19.6 | 1.001 | 92000 |
| $\alpha$ | 0.115 | 0.081 | 0.013 | 0.314 | 1.001 | 26000 |
| $\beta$ | 0.372 | 0.112 | 0.156 | 0.600 | 1.001 | 32000 |
| $r$ | 2.43 | 0.37 | 1.78 | 3.23 | 1.001 | 220000 |

##### 2003.2 infected

| Parameter | Mean | SD | 2.5% | 97.5% | $\hat{R}$ | $n_{\text{eff}}$ |
|---|---|---|---|---|---|---|
| $T_{\min}$ | 4.7 | 2.3 | 0.0 | 9.0 | 1.001 | 180000 |
| $T_{\text{opt}}$ | 13.3 | 3.1 | 5.8 | 17.7 | 1.001 | 29000 |
| $T_{\max}$ | 36.6 | 1.6 | 34.0 | 40.3 | 1.001 | 64000 |
| $\text{lf}_{\max}$ | 16.7 | 1.8 | 13.7 | 20.9 | 1.001 | 37000 |
| $\alpha$ | 0.265 | 0.101 | 0.056 | 0.442 | 1.001 | 14000 |
| $\beta$ | 0.368 | 0.063 | 0.255 | 0.502 | 1.001 | 22000 |
| $r$ | 2.25 | 0.30 | 1.72 | 2.89 | 1.001 | 250000 |

**2017.1 exposed uninfected**

| Parameter | Mean | SD | 2.5% | 97.5% | $\hat{R}$ | $n_{\text{eff}}$ |
|---|---|---|---|---|---|---|
| $T_{\text{min}}$ | 4.8 | 2.3 | 0.0 | 9.1 | 1.001 | 250000 |
| $T_{\text{opt}}$ | 14.3 | 5.1 | 4.2 | 22.6 | 1.001 | 190000 |
| $T_{\text{max}}$ | 35.9 | 1.8 | 33.3 | 40.0 | 1.001 | 250000 |
| $\text{lf}_{\text{max}}$ | 16.7 | 1.9 | 13.4 | 21.0 | 1.001 | 180000 |
| $\alpha$ | 0.305 | 0.168 | 0.023 | 0.614 | 1.001 | 210000 |
| $\beta$ | 0.458 | 0.134 | 0.251 | 0.781 | 1.001 | 250000 |
| $r$ | 1.69 | 0.28 | 1.19 | 2.30 | 1.001 | 160000 |

**2017.1 infected**

| Parameter | Mean | SD | 2.5% | 97.5% | $\hat{R}$ | $n_{\text{eff}}$ |
|---|---|---|---|---|---|---|
| $T_{\text{min}}$ | 9.3 | 1.1 | 6.0 | 10.0 | 1.001 | 22000 |
| $T_{\text{opt}}$ | 17.0 | 1.5 | 14.2 | 20.0 | 1.001 | 34000 |
| $T_{\text{max}}$ | 37.6 | 1.8 | 34.6 | 41.3 | 1.001 | 20000 |
| $\text{lf}_{\text{max}}$ | 19.3 | 1.8 | 16.1 | 23.3 | 1.001 | 47000 |
| $\alpha$ | 0.272 | 0.076 | 0.148 | 0.445 | 1.001 | 32000 |
| $\beta$ | 0.364 | 0.056 | 0.272 | 0.490 | 1.001 | 19000 |
| $r$ | 2.51 | 0.30 | 1.97 | 3.14 | 1.001 | 140000 |

#### Oviposition

We do not incorporate infection status effects in oviposition in the $\mathcal{R}_{0I}$ model. This is because, under standard assumptions of no vertical transmission of WNV, the only role of oviposition in transmission is through its effects on the total number of mosquitoes $M(T)$, where the traits of infected mosquitoes are not likely to have a strong effect due to the proportion of infected mosquitoes typically being low. However, we still measured oviposition in unexposed, exposed uninfected and infected mosquitoes. As for other unimodal traits, the proportion ovipositing was modeled with a flexTPC equation.

$$p_O = p_{O,\max}\left[\left(\frac{T - T_{\min}}{\alpha}\right)^{\alpha}\left(\frac{T_{\max} - T}{1-\alpha}\right)^{1-\alpha}\left(\frac{1}{T_{\max} - T_{\min}}\right)\right]^{\frac{\alpha(1-\alpha)}{\beta^2}}$$

where $p_{O,\max}$ is the maximum oviposition proportion (reached when $T = T_{\text{opt}}$), $T_{\min}$ and $T_{\max}$ the minimum and maximum temperatures, respectively, $\alpha \in (0,1)$ a parameter that determines the position of the thermal optimum $T_{\text{opt}}$ relative to the minimum and maximum (where $T_{\text{opt}} = \alpha T_{\max} + (1-\alpha)T_{\min}$) and $\beta$ a parameter that represents the approximate ratio of the thermal breadth at 88% maximum performance and the thermal tolerance range $T_{\max} - T_{\min}$. A separate curve was fit to mosquitoes unexposed to WNV, mosquitoes exposed to the 2003.2 or 2017.1 strain (but uninfected), and mosquitoes infected with the 2003.2 or 2017.1 strain.

##### Statistical model

Let $y_i$ denote the number of mosquitoes that laid eggs at temperature $T_i$ out of a total of $N_i$ mosquitoes. The likelihood can then be modeled by a binomial distribution.

$$y_i | N_i, T_i, \mathcal{P} \sim \text{Binomial}(p = p_O(T_i), N = N_i)$$

where $\mathcal{P} = \{T_{\text{min}}, T_{\text{max}}, p_{O,\text{max}}, \alpha, \beta\}$ is the parameters of the flexTPC model.

##### Prior distributions

$$\begin{aligned}
T_{\text{min}} &\sim \text{Normal}(\mu = 8.2°\text{C}, \sigma = 3°\text{C}) \\
T_{\text{max}} &\sim \text{Normal}(\mu = 33.2°\text{C}, \sigma = 3°\text{C}) \\
p_{O,\text{max}} &\sim \text{Uniform}(0, 1) \\
\alpha &\sim \text{Uniform}(0, 1) \\
\beta &\sim \text{Gamma}(\mu = 0.35, \sigma = 0.2)
\end{aligned}$$

The priors for the minimum and maximum temperatures are centered on previous estimates by *Shocket et al*, with an inflated standard deviation to account for possible differences between the mosquito populations in this work and those studied previously. The priors for $p_{O,\text{max}}$ and $\alpha$ are uniform priors that assume any maximum oviposition proportion or relative location of the thermal optimum is equally likely a priori. The prior for $\beta$ is weakly informative, preferring thermal breadths that are similar to common TPC models like the Briere or quadratic model but allowing other thermal breadths if necessary to describe the data.

##### Posterior distribution summaries

##### Unexposed

| Parameter | Mean | SD | 2.5% | 97.5% | $\hat{R}$ | $n_{\text{eff}}$ |
|---|---|---|---|---|---|---|
| $T_{\text{min}}$ | 11.2 | 1.1 | 9.7 | 13.9 | 1.001 | 250000 |
| $T_{\text{opt}}$ | 20.6 | 2.3 | 14.3 | 23.9 | 1.001 | 180000 |
| $T_{\text{max}}$ | 34.0 | 1.1 | 33.0 | 37.1 | 1.001 | 65000 |
| $p_{O,\text{max}}$ | 0.779 | 0.053 | 0.672 | 0.878 | 1.001 | 250000 |
| $\alpha$ | 0.411 | 0.114 | 0.113 | 0.588 | 1.001 | 110000 |
| $\beta$ | 0.487 | 0.106 | 0.309 | 0.721 | 1.001 | 250000 |

**2003.2 exposed uninfected**

| Parameter | Mean | SD | 2.5% | 97.5% | $\hat{R}$ | $n_{\text{eff}}$ |
|---|---|---|---|---|---|---|
| $T_{\text{min}}$ | 6.0 | 2.0 | 1.5 | 9.2 | 1.001 | 120000 |
| $T_{\text{opt}}$ | 28.2 | 2.0 | 24.2 | 31.7 | 1.001 | 250000 |
| $T_{\text{max}}$ | 31.8 | 0.9 | 30.2 | 33.0 | 1.001 | 150000 |
| $p_{O,\text{max}}$ | 0.583 | 0.109 | 0.379 | 0.800 | 1.001 | 250000 |
| $\alpha$ | 0.864 | 0.080 | 0.686 | 0.981 | 1.010 | 130000 |
| $\beta$ | 0.310 | 0.098 | 0.153 | 0.542 | 1.001 | 100000 |

**2003.2 infected**

| Parameter | Mean | SD | 2.5% | 97.5% | $\hat{R}$ | $n_{\text{eff}}$ |
|---|---|---|---|---|---|---|
| $T_{\text{min}}$ | 7.1 | 2.0 | 2.4 | 9.8 | 1.001 | 210000 |
| $T_{\text{opt}}$ | 25.0 | 2.2 | 21.0 | 30.1 | 1.001 | 210000 |
| $T_{\text{max}}$ | 31.7 | 0.8 | 30.1 | 33.0 | 1.001 | 250000 |
| $p_{O,\text{max}}$ | 0.643 | 0.061 | 0.526 | 0.763 | 1.001 | 250000 |
| $\alpha$ | 0.730 | 0.103 | 0.525 | 0.939 | 1.004 | 81000 |
| $\beta$ | 0.439 | 0.122 | 0.257 | 0.727 | 1.001 | 250000 |

**2017.1 exposed uninfected**

| Parameter | Mean | SD | 2.5% | 97.5% | $\hat{R}$ | $n_{\text{eff}}$ |
|---|---|---|---|---|---|---|
| $T_{\text{min}}$ | 10.7 | 2.0 | 5.5 | 14.0 | 1.001 | 170000 |
| $T_{\text{opt}}$ | 25.2 | 5.2 | 12.1 | 32.5 | 1.001 | 120000 |
| $T_{\text{max}}$ | 32.9 | 2.0 | 30.3 | 38.1 | 1.001 | 94000 |
| $p_{O,\text{max}}$ | 0.505 | 0.112 | 0.314 | 0.748 | 1.001 | 230000 |
| $\alpha$ | 0.662 | 0.263 | 0.026 | 0.989 | 1.001 | 120000 |
| $\beta$ | 0.475 | 0.185 | 0.206 | 0.925 | 1.001 | 200000 |

**2017.1 infected**

| Parameter | Mean | SD | 2.5% | 97.5% | $\hat{R}$ | $n_{\text{eff}}$ |
|---|---|---|---|---|---|---|
| $T_{\text{min}}$ | 10.2 | 1.9 | 5.6 | 13.6 | 1.001 | 250000 |
| $T_{\text{opt}}$ | 24.6 | 1.1 | 22.2 | 26.5 | 1.001 | 160000 |
| $T_{\text{max}}$ | 33.5 | 0.7 | 33.0 | 35.4 | 1.001 | 48000 |
| $p_{O,\text{max}}$ | 0.766 | 0.053 | 0.658 | 0.866 | 1.001 | 250000 |
| $\alpha$ | 0.614 | 0.069 | 0.465 | 0.733 | 1.001 | 130000 |
| $\beta$ | 0.353 | 0.084 | 0.218 | 0.545 | 1.001 | 220000 |

#### Relative $\mathcal{R}_0$

To evaluate the effect of incorporating infection-dependent mosquito life-history traits on WNV transmission, we compare the predictions of a relative $\mathcal{R}_0$ based only on unexposed mosquito traits

$$\text{relative } \mathcal{R}_0 = \sqrt{\frac{a^2(T)bc(T)e^{-\frac{\mu(T)}{\mathrm{PDR}(T)}}M(T)}{\mu(T)}}$$

as is currently standard in the literature, with an approach derived here that incorporates traits of infected mosquitoes in processes that involve them (see the Temperature-dependent relative $\mathcal{R}_0$ section above for the derivation).

$$\text{relative } \mathcal{R}_0 = \sqrt{\frac{a(T)a_I(T)bc(T)e^{-\frac{\mu_I(T)}{\mathrm{PDR}(T)}}M(T)}{\mu_I(T)}}$$

In either case, mosquito abundance was estimated as

$$M(T) = \frac{a_G(T)\mathrm{EFGC}(T)\mathrm{EV}(T)p_{LA}(T)\mathrm{MDR}(T)}{\mu^2(T)}$$

as described above.

These expressions are calculated from various mosquito life-history traits, as described earlier in the Temperature-dependent $\mathcal{R}_0$ section. We used vestimates from the TPC fits and data presented here for mosquito biting rates $a(T)$, $a_I(T)$, death rates $\mu(T)$, $\mu_I(T)$ (calculated as the inverse of lifespan) and vector competence $bc(T)$, with the estimates for $a_I(T), \mu_I(T)$ and $bc(T)$ being viral strain specific. For traits not measured here, we used TPCs previously estimated for *Culex pipiens* and WNV in [1].

| Transmission model trais | Strain | $T_{\min}$ | | | $T_{\mathrm{opt}}$ | | | $T_{\max}$ | | |
|---|---|---|---|---|---|---|---|---|---|---|
| | | Mean | 2.5% | 97.5% | Mean | 2.5% | 97.5% | Mean | 2.5% | 97.5% |
| unexposed | 2003.2 | 12.6 | 9.3 | 15.0 | 23.8 | 22.9 | 24.9 | 33.4 | 30.8 | 34.7 |
| unexposed | 2017.1 | 12.4 | 9.2 | 15.0 | 23.9 | 22.9 | 25.0 | 34.1 | 30.8 | 37.3 |
| unexposed+infected | 2003.2 | 13.3 | 9.9 | 16.7 | 23.9 | 22.9 | 25.1 | 32.6 | 29.1 | 34.5 |
| unexposed+infected | 2017.1 | 12.6 | 9.8 | 15.0 | 24.3 | 23.4 | 25.2 | 33.5 | 30.7 | 36.4 |
