## Supplemental Figures and Tables for "Mosquito infection-mediated trait variation alters temperature-dependent transmission of West Nile virus"

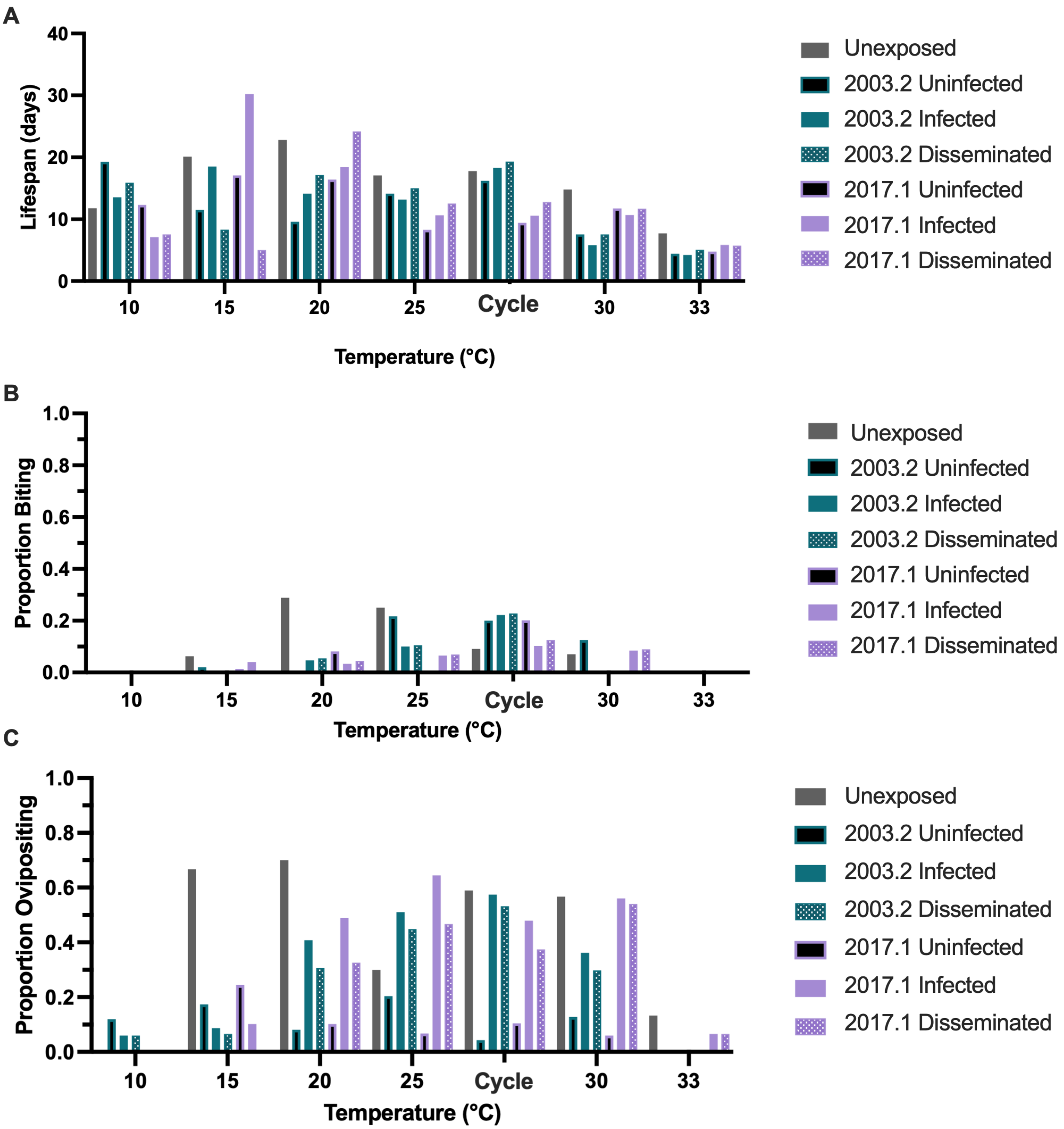

**Figure S1. Life history traits of 2003.2 and 2017.1 exposed *Culex pipiens* of various infection status and unexposed *Culex pipiens* across temperature.** Unexposed in gray, 2003.2 exposed mosquitoes are depicted as uninfected in black with a blue outline, infected in solid blue, and disseminated in the pattern. 2017.1 exposed mosquitoes are depicted as uninfected in black with a purple outline, infected in solid purple, and disseminated in the purple pattern. (**A**) Mean longevity; (**B**) Proportion biting; (**C**) Proportion of mosquitoes laying egg rafts.

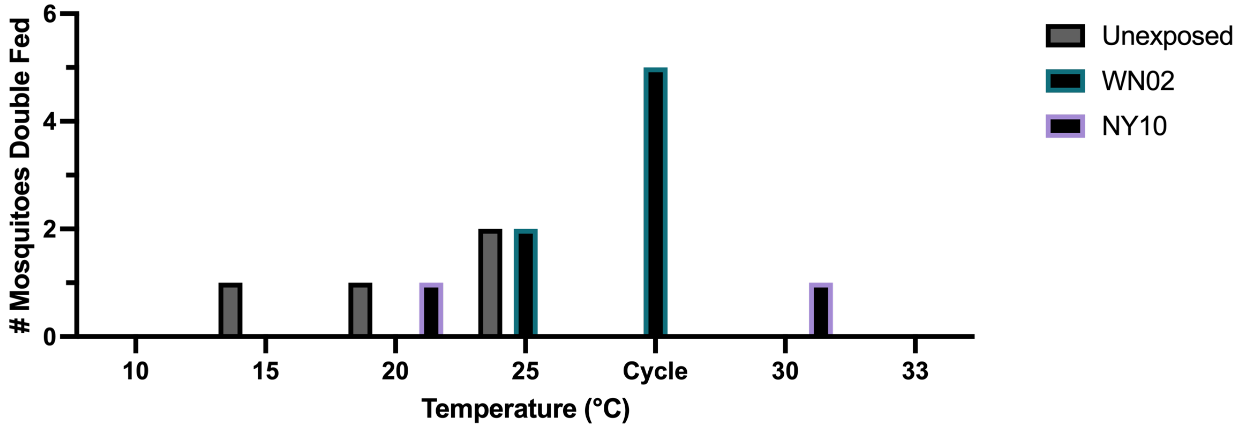

**Figure S2. The total number of mosquitoes that acquired more than one blood meal course of the study.** Unexposed in gray, 2003.2 exposed mosquitoes are depicted as uninfected in black with a blue outline and 2017.1 exposed mosquitoes are depicted as uninfected in black with a purple outline.

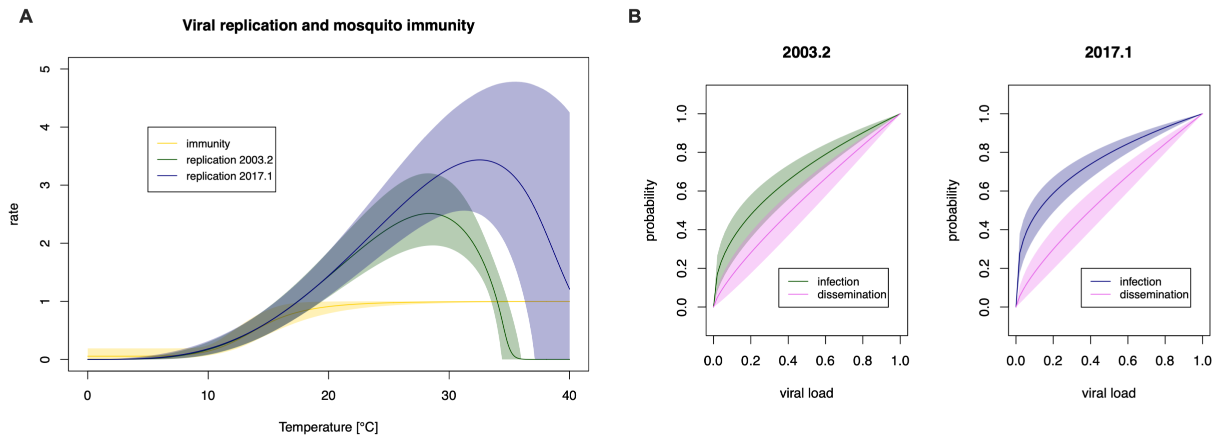

**Figure S3. Estimated latent TPCs giving rise to bimodal vector competence curve.** We developed a novel semi-mechanistic model that can describe bimodal TPCs (see Appendix for model derivation, parameterization and inference). The model was fit to the vector competence data (all panels of Fig. 3). Latent TPCs for viral replication and viral immunity were inferred from the data. **A)** Our model assumes that there is a tradeoff between unimodal viral replication (blue and green curves) and monotonic mosquito immunity (yellow curve). This tradeoff determines an initial viral load at the initial stages of infection. **B)** The probability of a mosquito developing an infection (green, blue) or a disseminated infection (pink) is assumed to increase monotonically with this initial viral load. Lines correspond to mean posterior estimates and shaded regions to 95% credible intervals.

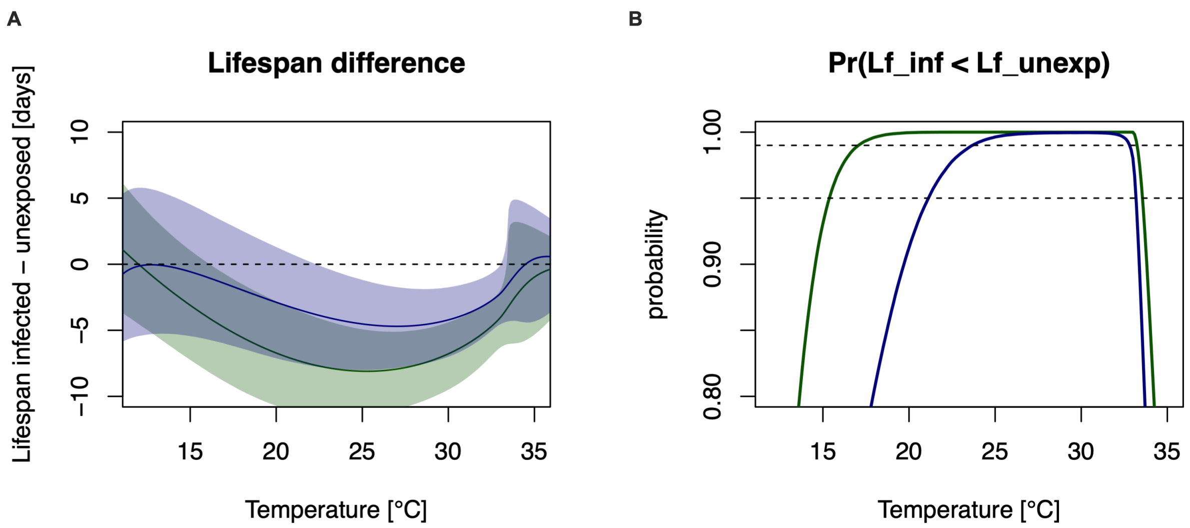

**Figure S4. WNV infection decreases mosquito lifespan in a wide temperature range. A)** Estimated difference between the lifespan of mosquitoes infected with the 2003.2 (green) or 2017.2 (purple) strains, and unexposed mosquitoes at each temperature. Lines correspond to posterior means and shaded regions to 95% credible intervals. **B)** Posterior probability that the lifespan of the infected mosquitoes is lower than for the unexposed mosquitoes for each strain at each temperature. 95% and 99% posterior probabilities are shown as dotted lines.

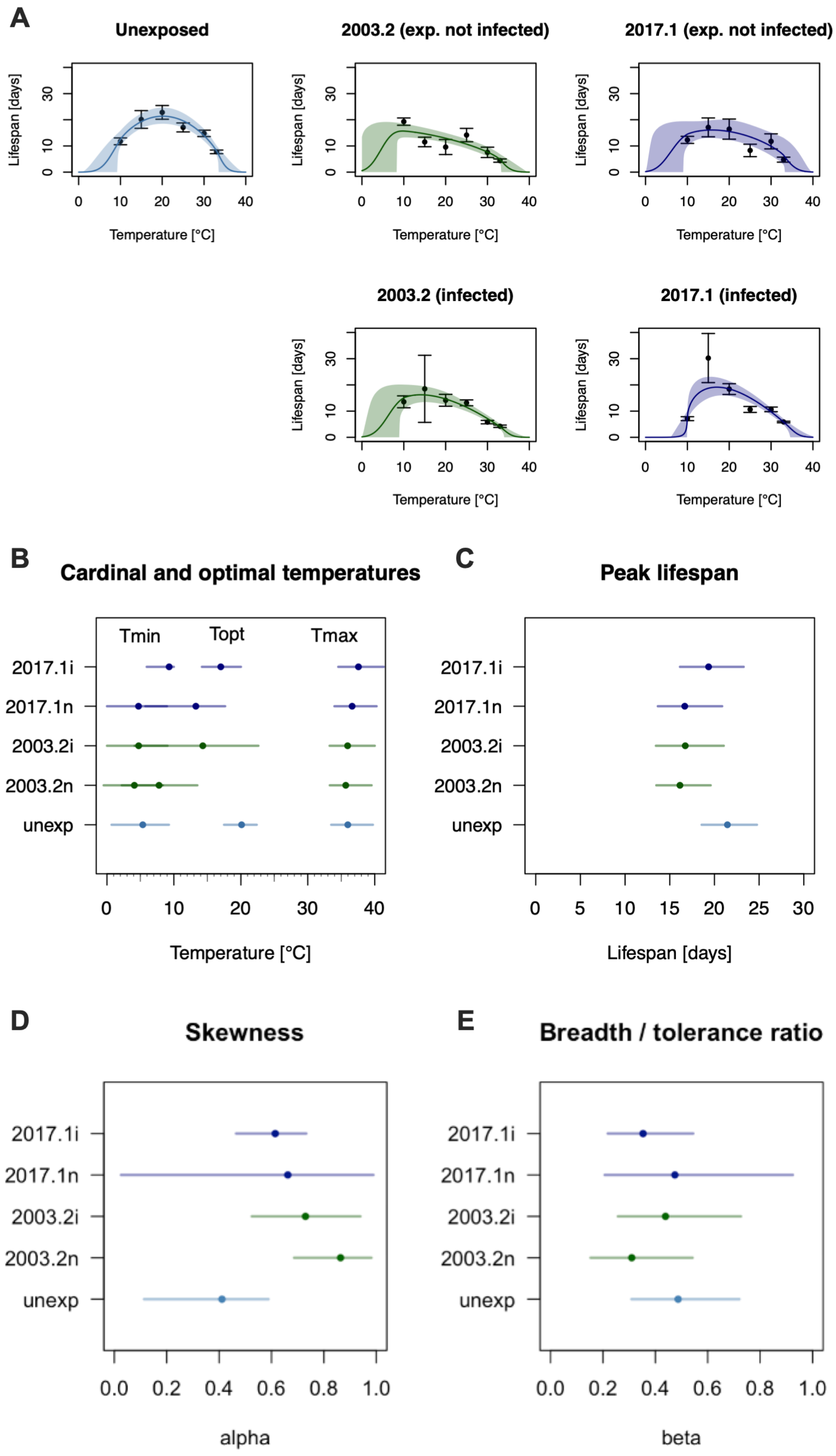
A
Unexposed
2003.2 (exp. not infected)
2017.1 (exp. not infected)
2003.2 (infected)
2017.1 (infected)
Lifespan [days]
Temperature [°C]
B
Cardinal and optimal temperatures
Tmin
Topt
Tmax
2017.1i
2017.1n
2003.2i
2003.2n
unexp
Temperature [°C]
C
Peak lifespan
Lifespan [days]
D
Skewness
alpha
E
Breadth / tolerance ratio
beta

**Figure S5. Thermal performance curves and estimated parameters for lifespan. A)** Individual thermal performance curves for each group of unexposed, exposed Uninfected and infected groups for the 2003.2 and 2017.1 strains. Data are shown as points and standard errors as error bars. Bayesian flexTPC model fits are shown as lines representing posterior means and shaded regions representing 95% credible intervals. **B)** Estimated parameters from the flexTPC model (minimum, optimum and maximum temperatures. **C)** Peak lifespan. **D)** Curve skewness. **E)** Thermal breadth/tolerance ratio) for each group (unexposed, and exposed uninfected (n) and infected (i) mosquitoes for the 2003.2 and 2017.2 WNV strains).

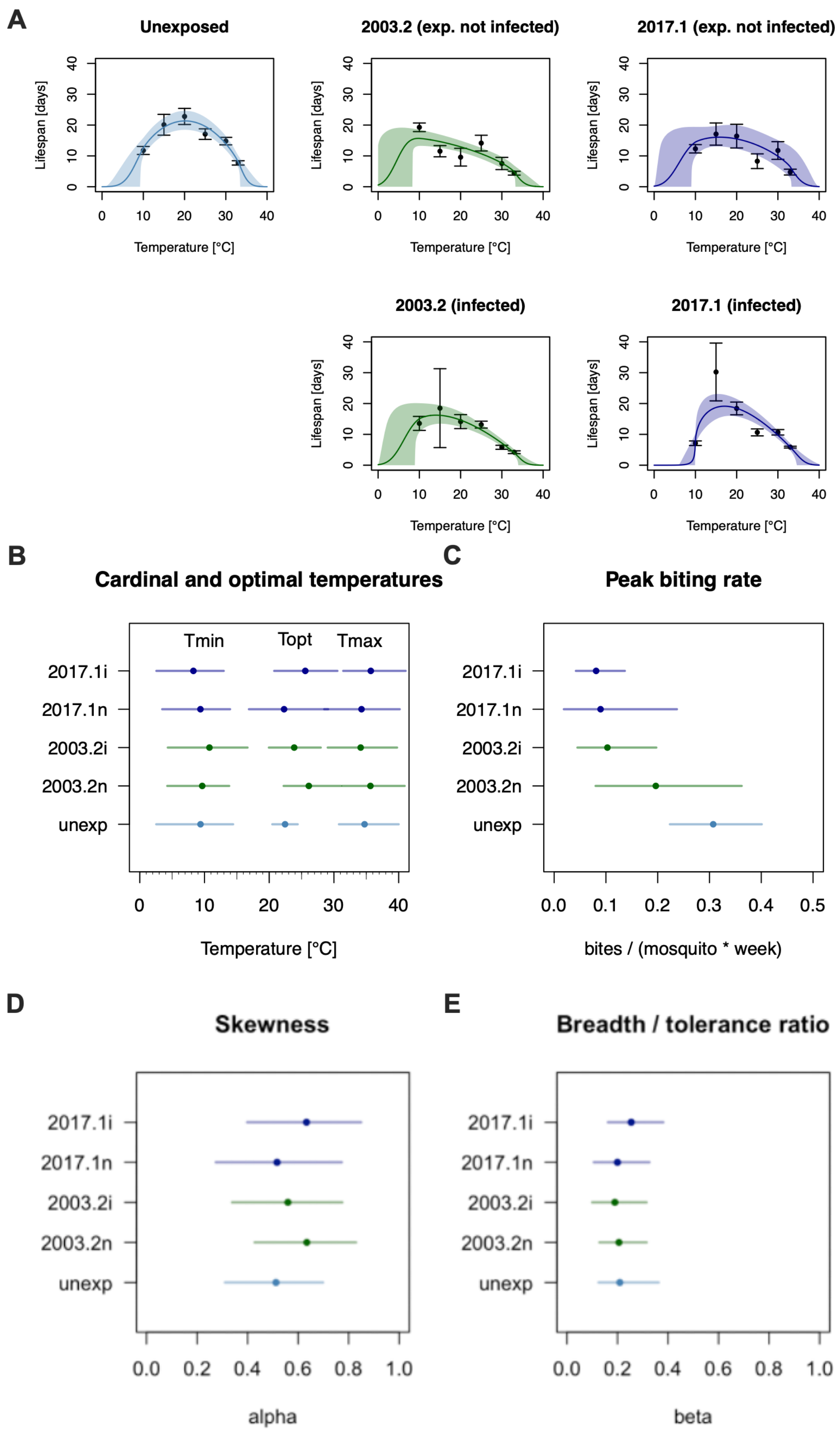
A
Unexposed
2003.2 (exp. not infected)
2017.1 (exp. not infected)
2003.2 (infected)
2017.1 (infected)
Lifespan [days]
Temperature [°C]
B
Cardinal and optimal temperatures
Tmin
Topt
Tmax
2017.1i
2017.1n
2003.2i
2003.2n
unexp
Temperature [°C]
C
Peak biting rate
bites / (mosquito * week)
D
Skewness
alpha
E
Breadth / tolerance ratio
beta

**Figure S6. Thermal performance curves and estimated parameters for biting rate. A)** Individual thermal performance curves for each group of unexposed, exposed Uninfected and infected groups for the 2003.2 and 2017.1 strains. Data are shown as points and standard errors as error bars. Bayesian flexTPC model fits are shown as lines representing posterior means and shaded regions representing 95% credible intervals. **B)** Estimated parameters from the flexTPC model (minimum, optimum and maximum temperatures) **C)** Peak biting rate. **D)** curve skewness. **E)** thermal breadth/tolerance ratio) for each group (unexposed, and exposed uninfected (n) and infected (i) mosquitoes for the 2003.2 and 2017.2 WNV strains).

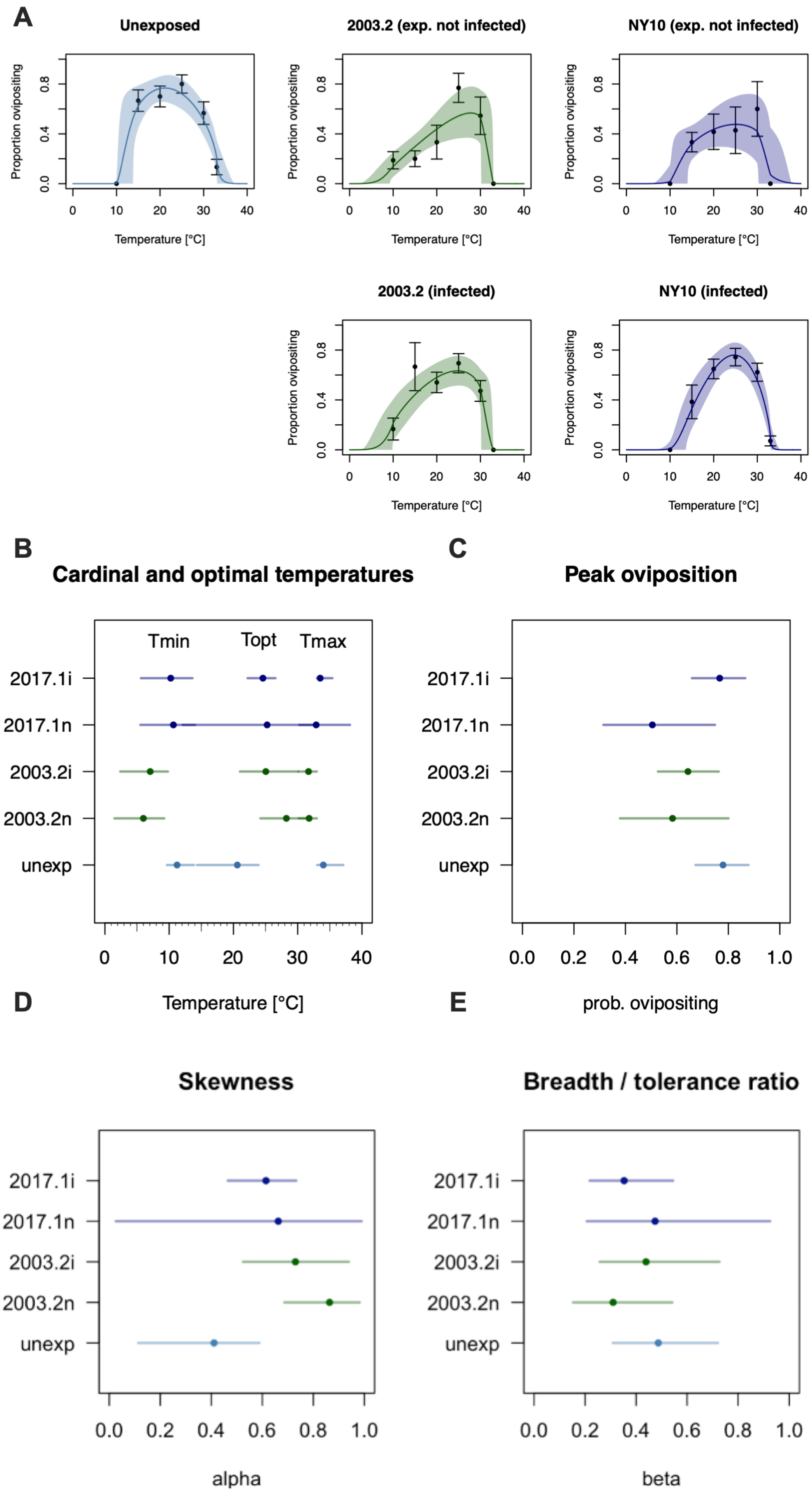
A
Unexposed
2003.2 (exp. not infected)
NY10 (exp. not infected)
2003.2 (infected)
NY10 (infected)
Proportion ovipositing
Temperature [°C]
B
Cardinal and optimal temperatures
Tmin
Topt
Tmax
2017.1i
2017.1n
2003.2i
2003.2n
unexp
Temperature [°C]
C
Peak oviposition
prob. ovipositing
D
Skewness
alpha
E
Breadth / tolerance ratio
beta

**Figure S7. Thermal performance curves and estimated parameters for oviposition. A)** Individual thermal performance curves for each group of unexposed, exposed Uninfected and infected groups for the 2003.2 and 2017.1 strains. Data are shown as points and standard errors as error bars. Bayesian flexTPC model fits are shown as lines representing posterior means and shaded regions representing 95% credible intervals. **B)** Estimated parameters from the flexTPC model (minimum, optimum and maximum temperatures) **C)** Peak oviposition **D)** Curve skewness. **E)** Thermal breadth/tolerance ratio) for each group (unexposed, and exposed uninfected (n) and infected (i) mosquitoes for the 2003.2 and 2017.2 WNV strains).

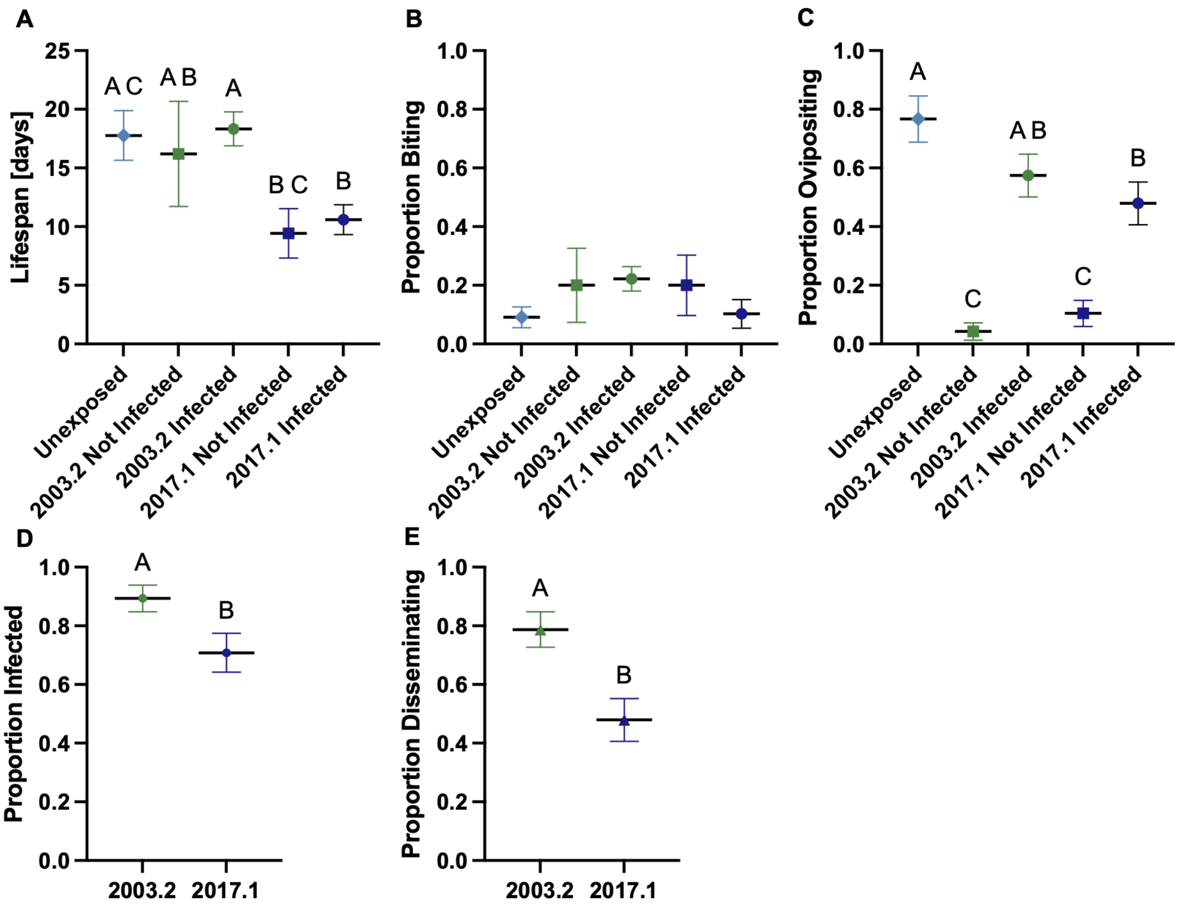

**Figure S8. *Cx. pipiens* life history traits at cycling 25°C**. (**A**) Adult lifespan of *Cx. pipiens* mosquitoes by infection status. (**B**) Proportion of mosquitoes blood-feeding by infection status. (**C**) Proportion of mosquitoes ovipositing by infection status. For panels A–C, statistically significant differences are indicated by different letters ($p \leq 0.05$; one-way ANOVA with Tukey's multiple comparisons post hoc test). (**D**) Proportion infected. (**E**) Proportion disseminated. For panels D–E, statistically significant differences are indicated by different letters ($p \leq 0.05$; Welch's t-test). All panels show mean ± SEM.

**Table S1. Pairwise differences in lifespan among infection statuses within each temperature were evaluated using Two-way ANOVA with interaction, followed by Tukey-adjusted post hoc.**

| Temperature | Pairwise | Magnitude | Direction | P-value | Significance |
|---|---|---|---|---|---|
| 10 | Unexposed - Disseminated 2017.1 | 4.228 | Positive | 0.553 | No |
| 10 | Unexposed - Infected 2017.1 | 4.638 | Positive | 0.148 | No |
| 10 | Unexposed - Not Infected 2017.1 | 0.546 | Negative | 1.000 | No |
| 10 | Unexposed - Disseminated 2003.2 | 4.150 | Negative | 0.608 | No |
| 10 | Unexposed - Infected 2003.2 | 1.789 | Negative | 0.980 | No |
| 10 | Unexposed - Not Infected 2003.2 | 7.515 | Negative | 0.001 | Yes |
| 10 | Disseminated 2017.1 - Infected 2017.1 | 0.409 | Positive | 1.000 | No |
| 10 | Disseminated 2017.1 - Not Infected 2017.1 | 4.774 | Negative | 0.548 | No |
| 10 | Disseminated 2017.1 - Disseminated 2003.2 | 8.378 | Negative | 0.056 | No |
| 10 | Disseminated 2017.1 - Infected 2003.2 | 6.017 | Negative | 0.236 | No |
| 10 | Disseminated 2017.1 - Not Infected 2003.2 | 11.743 | Negative | 0.000 | Yes |
| 10 | Infected 2017.1 - Not Infected 2017.1 | 5.183 | Negative | 0.217 | No |
| 10 | Infected 2017.1 - Disseminated 2003.2 | 8.788 | Negative | 0.007 | Yes |
| 10 | Infected 2017.1 - Infected 2003.2 | 6.427 | Negative | 0.041 | Yes |
| 10 | Infected 2017.1 - Not Infected 2003.2 | 12.152 | Negative | 0.000 | Yes |
| 10 | Not Infected 2017.1 - Disseminated 2003.2 | 3.604 | Negative | 0.836 | No |
| 10 | Not Infected 2017.1 - Infected 2003.2 | 1.243 | Negative | 0.999 | No |
| 10 | Not Infected 2017.1 - Not Infected 2003.2 | 6.969 | Negative | 0.027 | Yes |
| 10 | Disseminated 2003.2 - Infected 2003.2 | 2.361 | Positive | 0.973 | No |
| 10 | Disseminated 2003.2 - Not Infected 2003.2 | 3.365 | Negative | 0.800 | No |
| 10 | Infected 2003.2 - Not Infected 2003.2 | 5.726 | Negative | 0.096 | No |
| 15 | Unexposed - Disseminated 2017.1 | 15.100 | Positive | 0.947 | No |
| 15 | Unexposed - Infected 2017.1 | 10.131 | Negative | 0.736 | No |
| 15 | Unexposed - Not Infected 2017.1 | 3.017 | Positive | 0.997 | No |
| 15 | Unexposed - Disseminated 2003.2 | 11.767 | Positive | 0.961 | No |
| 15 | Unexposed - Infected 2003.2 | 1.600 | Positive | 1.000 | No |
| 15 | Unexposed - Not Infected 2003.2 | 8.575 | Positive | 0.576 | No |
| 15 | Disseminated 2017.1 - Infected 2017.1 | 25.231 | Negative | 0.651 | No |
| 15 | Disseminated 2017.1 - Not Infected 2017.1 | 12.083 | Negative | 0.982 | No |
| 15 | Disseminated 2017.1 - Disseminated 2003.2 | 3.333 | Negative | 1.000 | No |
| 15 | Disseminated 2017.1 - Infected 2003.2 | 13.500 | Negative | 0.982 | No |
| 15 | Disseminated 2017.1 - Not Infected 2003.2 | 6.525 | Negative | 0.999 | No |
| 15 | Infected 2017.1 - Not Infected 2017.1 | 13.147 | Positive | 0.410 | No |
| 15 | Infected 2017.1 - Disseminated 2003.2 | 21.897 | Positive | 0.619 | No |
| 15 | Infected 2017.1 - Infected 2003.2 | 11.731 | Positive | 0.901 | No |
| 15 | Infected 2017.1 - Not Infected 2003.2 | 18.706 | Positive | 0.064 | No |
| 15 | Not Infected 2017.1 - Disseminated 2003.2 | 8.750 | Positive | 0.991 | No |
| 15 | Not Infected 2017.1 - Infected 2003.2 | 1.417 | Negative | 1.000 | No |
| 15 | Not Infected 2017.1 - Not Infected 2003.2 | 5.558 | Positive | 0.893 | No |
| 15 | Disseminated 2003.2 - Infected 2003.2 | 10.167 | Negative | 0.992 | No |
| 15 | Disseminated 2003.2 - Not Infected 2003.2 | 3.192 | Negative | 1.000 | No |

| | | | | | |
|---|---|---|---|---|---|
| 15 | Infected 2003.2 - Not Infected 2003.2 | 6.975 | Positive | 0.985 | No |
| 20 | Unexposed - Disseminated 2017.1 | 1.390 | Negative | 1.000 | No |
| 20 | Unexposed - Infected 2017.1 | 4.395 | Positive | 0.840 | No |
| 20 | Unexposed - Not Infected 2017.1 | 6.383 | Positive | 0.810 | No |
| 20 | Unexposed - Disseminated 2003.2 | 5.640 | Positive | 0.719 | No |
| 20 | Unexposed - Infected 2003.2 | 8.665 | Positive | 0.129 | No |
| 20 | Unexposed - Not Infected 2003.2 | 13.217 | Positive | 0.069 | No |
| 20 | Disseminated 2017.1 - Infected 2017.1 | 5.785 | Positive | 0.703 | No |
| 20 | Disseminated 2017.1 - Not Infected 2017.1 | 7.774 | Positive | 0.688 | No |
| 20 | Disseminated 2017.1 - Disseminated 2003.2 | 7.030 | Positive | 0.578 | No |
| 20 | Disseminated 2017.1 - Infected 2003.2 | 10.055 | Positive | 0.098 | No |
| 20 | Disseminated 2017.1 - Not Infected 2003.2 | 14.607 | Positive | 0.049 | Yes |
| 20 | Infected 2017.1 - Not Infected 2017.1 | 1.989 | Positive | 0.999 | No |
| 20 | Infected 2017.1 - Disseminated 2003.2 | 1.245 | Positive | 1.000 | No |
| 20 | Infected 2017.1 - Infected 2003.2 | 4.270 | Positive | 0.822 | No |
| 20 | Infected 2017.1 - Not Infected 2003.2 | 8.822 | Positive | 0.440 | No |
| 20 | Not Infected 2017.1 - Disseminated 2003.2 | 0.743 | Negative | 1.000 | No |
| 20 | Not Infected 2017.1 - Infected 2003.2 | 2.282 | Positive | 0.999 | No |
| 20 | Not Infected 2017.1 - Not Infected 2003.2 | 6.833 | Positive | 0.878 | No |
| 20 | Disseminated 2003.2 - Infected 2003.2 | 3.025 | Positive | 0.977 | No |
| 20 | Disseminated 2003.2 - Not Infected 2003.2 | 7.577 | Positive | 0.684 | No |
| 20 | Infected 2003.2 - Not Infected 2003.2 | 4.552 | Positive | 0.950 | No |
| 25 | Unexposed - Disseminated 2017.1 | 4.515 | Positive | 0.229 | No |
| 25 | Unexposed - Infected 2017.1 | 6.426 | Positive | 0.008 | Yes |
| 25 | Unexposed - Not Infected 2017.1 | 8.781 | Positive | 0.075 | No |
| 25 | Unexposed - Disseminated 2003.2 | 2.032 | Positive | 0.940 | No |
| 25 | Unexposed - Infected 2003.2 | 3.900 | Positive | 0.337 | No |
| 25 | Unexposed - Not Infected 2003.2 | 2.913 | Positive | 0.898 | No |
| 25 | Disseminated 2017.1 - Infected 2017.1 | 1.911 | Positive | 0.940 | No |
| 25 | Disseminated 2017.1 - Not Infected 2017.1 | 4.266 | Positive | 0.817 | No |
| 25 | Disseminated 2017.1 - Disseminated 2003.2 | 2.483 | Negative | 0.861 | No |
| 25 | Disseminated 2017.1 - Infected 2003.2 | 0.615 | Negative | 1.000 | No |
| 25 | Disseminated 2017.1 - Not Infected 2003.2 | 1.602 | Negative | 0.995 | No |
| 25 | Infected 2017.1 - Not Infected 2017.1 | 2.355 | Positive | 0.987 | No |
| 25 | Infected 2017.1 - Disseminated 2003.2 | 4.393 | Negative | 0.195 | No |
| 25 | Infected 2017.1 - Infected 2003.2 | 2.526 | Negative | 0.757 | No |
| 25 | Infected 2017.1 - Not Infected 2003.2 | 3.513 | Negative | 0.754 | No |
| 25 | Not Infected 2017.1 - Disseminated 2003.2 | 6.749 | Negative | 0.318 | No |
| 25 | Not Infected 2017.1 - Infected 2003.2 | 4.881 | Negative | 0.683 | No |
| 25 | Not Infected 2017.1 - Not Infected 2003.2 | 5.868 | Negative | 0.621 | No |
| 25 | Disseminated 2003.2 - Infected 2003.2 | 1.868 | Positive | 0.951 | No |
| 25 | Disseminated 2003.2 - Not Infected 2003.2 | 0.881 | Positive | 1.000 | No |
| 25 | Infected 2003.2 - Not Infected 2003.2 | 0.987 | Negative | 1.000 | No |
| 30 | Unexposed - Disseminated 2017.1 | 3.082 | Positive | 0.249 | No |

| | | | | | |
|---|---|---|---|---|---|
| 30 | Unexposed - Infected 2017.1 | 4.126 | Positive | 0.028 | Yes |
| 30 | Unexposed - Not Infected 2017.1 | 3.050 | Positive | 0.944 | No |
| 30 | Unexposed - Disseminated 2003.2 | 7.235 | Positive | 0.000 | Yes |
| 30 | Unexposed - Infected 2003.2 | 8.994 | Positive | 0.000 | Yes |
| 30 | Unexposed - Not Infected 2003.2 | 7.255 | Positive | 0.005 | Yes |
| 30 | Disseminated 2017.1 - Infected 2017.1 | 1.044 | Positive | 0.977 | No |
| 30 | Disseminated 2017.1 - Not Infected 2017.1 | 0.032 | Negative | 1.000 | No |
| 30 | Disseminated 2017.1 - Disseminated 2003.2 | 4.153 | Positive | 0.069 | No |
| 30 | Disseminated 2017.1 - Infected 2003.2 | 5.912 | Positive | 0.000 | Yes |
| 30 | Disseminated 2017.1 - Not Infected 2003.2 | 4.172 | Positive | 0.292 | No |
| 30 | Infected 2017.1 - Not Infected 2017.1 | 1.076 | Negative | 1.000 | No |
| 30 | Infected 2017.1 - Disseminated 2003.2 | 3.109 | Positive | 0.297 | No |
| 30 | Infected 2017.1 - Infected 2003.2 | 4.868 | Positive | 0.002 | Yes |
| 30 | Infected 2017.1 - Not Infected 2003.2 | 3.128 | Positive | 0.624 | No |
| 30 | Not Infected 2017.1 - Disseminated 2003.2 | 4.185 | Positive | 0.801 | No |
| 30 | Not Infected 2017.1 - Infected 2003.2 | 5.944 | Positive | 0.390 | No |
| 30 | Not Infected 2017.1 - Not Infected 2003.2 | 4.205 | Positive | 0.849 | No |
| 30 | Disseminated 2003.2 - Infected 2003.2 | 1.760 | Positive | 0.895 | No |
| 30 | Disseminated 2003.2 - Not Infected 2003.2 | 0.020 | Positive | 1.000 | No |
| 30 | Infected 2003.2 - Not Infected 2003.2 | 1.740 | Negative | 0.970 | No |
| 33 | Unexposed - Disseminated 2017.1 | 2.004 | Positive | 0.026 | Yes |
| 33 | Unexposed - Infected 2017.1 | 1.876 | Positive | 0.037 | Yes |
| 33 | Unexposed - Not Infected 2017.1 | 2.983 | Positive | 0.298 | No |
| 33 | Unexposed - Disseminated 2003.2 | 2.683 | Positive | 0.006 | Yes |
| 33 | Unexposed - Infected 2003.2 | 3.521 | Positive | 0.000 | Yes |
| 33 | Unexposed - Not Infected 2003.2 | 3.333 | Positive | 0.001 | Yes |
| 33 | Disseminated 2017.1 - Infected 2017.1 | 0.127 | Negative | 1.000 | No |
| 33 | Disseminated 2017.1 - Not Infected 2017.1 | 0.980 | Positive | 0.990 | No |
| 33 | Disseminated 2017.1 - Disseminated 2003.2 | 0.680 | Positive | 0.961 | No |
| 33 | Disseminated 2017.1 - Infected 2003.2 | 1.518 | Positive | 0.168 | No |
| 33 | Disseminated 2017.1 - Not Infected 2003.2 | 1.330 | Positive | 0.610 | No |
| 33 | Infected 2017.1 - Not Infected 2017.1 | 1.107 | Positive | 0.981 | No |
| 33 | Infected 2017.1 - Disseminated 2003.2 | 0.807 | Positive | 0.905 | No |
| 33 | Infected 2017.1 - Infected 2003.2 | 1.645 | Positive | 0.085 | No |
| 33 | Infected 2017.1 - Not Infected 2003.2 | 1.457 | Positive | 0.479 | No |
| 33 | Not Infected 2017.1 - Disseminated 2003.2 | 0.300 | Negative | 1.000 | No |
| 33 | Not Infected 2017.1 - Infected 2003.2 | 0.538 | Positive | 1.000 | No |
| 33 | Not Infected 2017.1 - Not Infected 2003.2 | 0.350 | Positive | 1.000 | No |
| 33 | Disseminated 2003.2 - Infected 2003.2 | 0.838 | Positive | 0.907 | No |
| 33 | Disseminated 2003.2 - Not Infected 2003.2 | 0.650 | Positive | 0.989 | No |
| 33 | Infected 2003.2 - Not Infected 2003.2 | 0.188 | Negative | 1.000 | No |
| Cycle | Unexposed - Disseminated 2017.1 | 4.984 | Positive | 0.420 | No |
| Cycle | Unexposed - Infected 2017.1 | 7.178 | Positive | 0.028 | Yes |
| Cycle | Unexposed - Not Infected 2017.1 | 8.338 | Positive | 0.069 | No |
| Cycle | Unexposed - Disseminated 2003.2 | 1.558 | Negative | 0.992 | No |

| | | | | | |
|---|---|---|---|---|---|
| Cycle | Unexposed - Infected 2003.2 | 0.567 | Negative | 1.000 | No |
| Cycle | Unexposed - Not Infected 2003.2 | 1.567 | Positive | 1.000 | No |
| Cycle | Disseminated 2017.1 - Infected 2017.1 | 2.194 | Positive | 0.972 | No |
| Cycle | Disseminated 2017.1 - Not Infected 2017.1 | 3.354 | Positive | 0.928 | No |
| Cycle | Disseminated 2017.1 - Disseminated 2003.2 | 6.542 | Negative | 0.095 | No |
| Cycle | Disseminated 2017.1 - Infected 2003.2 | 5.551 | Negative | 0.214 | No |
| Cycle | Disseminated 2017.1 - Not Infected 2003.2 | 3.417 | Negative | 0.988 | No |
| Cycle | Infected 2017.1 - Not Infected 2017.1 | 1.160 | Positive | 1.000 | No |
| Cycle | Infected 2017.1 - Disseminated 2003.2 | 8.736 | Negative | 0.001 | Yes |
| Cycle | Infected 2017.1 - Infected 2003.2 | 7.745 | Negative | 0.005 | Yes |
| Cycle | Infected 2017.1 - Not Infected 2003.2 | 5.612 | Negative | 0.850 | No |
| Cycle | Not Infected 2017.1 - Disseminated 2003.2 | 9.896 | Negative | 0.010 | Yes |
| Cycle | Not Infected 2017.1 - Infected 2003.2 | 8.905 | Negative | 0.026 | Yes |
| Cycle | Not Infected 2017.1 - Not Infected 2003.2 | 6.771 | Negative | 0.777 | No |
| Cycle | Disseminated 2003.2 - Infected 2003.2 | 0.991 | Positive | 0.999 | No |
| Cycle | Disseminated 2003.2 - Not Infected 2003.2 | 3.124 | Positive | 0.991 | No |
| Cycle | Infected 2003.2 - Not Infected 2003.2 | 2.133 | Positive | 0.999 | No |

**Table S2. Pairwise differences in blood-feeding proportions among infection statuses within each temperature were evaluated using Fisher's exact tests with Bonferroni correction for multiple comparisons.**

| Temperature | Pairwise | Magnitude | Direction | P-value | Significance |
|---|---|---|---|---|---|
| 10 | Unexposed vs 2003.2_Uninfected | 0.000 | Negative | 1.000 | No |
| 10 | Unexposed vs 2003.2_Infected | 0.000 | Negative | 1.000 | No |
| 10 | Unexposed vs 2003.2_Dissemination | 0.000 | Negative | 1.000 | No |
| 10 | Unexposed vs 2017.1_Uninfected | 0.000 | Negative | 1.000 | No |
| 10 | Unexposed vs 2017.1_Infected | 0.000 | Negative | 1.000 | No |
| 10 | Unexposed vs 2017.1_Disseminated | 0.000 | Negative | 1.000 | No |
| 10 | 2003.2_Uninfected vs 2003.2_Infected | 0.000 | Negative | 1.000 | No |
| 10 | 2003.2_Uninfected vs 2003.2_Dissemination | 0.000 | Negative | 1.000 | No |
| 10 | 2003.2_Uninfected vs 2017.1_Uninfected | 0.000 | Negative | 1.000 | No |
| 10 | 2003.2_Uninfected vs 2017.1_Infected | 0.000 | Negative | 1.000 | No |
| 10 | 2003.2_Uninfected vs 2017.1_Disseminated | 0.000 | Negative | 1.000 | No |
| 10 | 2003.2_Infected vs 2003.2_Dissemination | 0.000 | Negative | 1.000 | No |
| 10 | 2003.2_Infected vs 2017.1_Uninfected | 0.000 | Negative | 1.000 | No |
| 10 | 2003.2_Infected vs 2017.1_Infected | 0.000 | Negative | 1.000 | No |
| 10 | 2003.2_Infected vs 2017.1_Disseminated | 0.000 | Negative | 1.000 | No |
| 10 | 2003.2_Dissemination vs 2017.1_Uninfected | 0.000 | Negative | 1.000 | No |
| 10 | 2003.2_Dissemination vs 2017.1_Infected | 0.000 | Negative | 1.000 | No |
| 10 | 2003.2_Dissemination vs 2017.1_Disseminated | 0.000 | Negative | 1.000 | No |
| 10 | 2017.1_Uninfected vs 2017.1_Infected | 0.000 | Negative | 1.000 | No |
| 10 | 2017.1_Uninfected vs 2017.1_Disseminated | 0.000 | Negative | 1.000 | No |
| 10 | 2017.1_Infected vs 2017.1_Disseminated | 0.000 | Negative | 1.000 | No |
| 15 | Unexposed vs 2003.2_Uninfected | 0.043 | Positive | 1.000 | No |
| 15 | Unexposed vs 2003.2_Infected | 0.063 | Positive | 1.000 | No |
| 15 | Unexposed vs 2003.2_Dissemination | 0.063 | Positive | 1.000 | No |
| 15 | Unexposed vs 2017.1_Uninfected | 0.049 | Positive | 1.000 | No |
| 15 | Unexposed vs 2017.1_Infected | 0.023 | Positive | 1.000 | No |
| 15 | Unexposed vs 2017.1_Disseminated | | | 1.000 | No |
| 15 | 2003.2_Uninfected vs 2003.2_Infected | 0.020 | Positive | 1.000 | No |
| 15 | 2003.2_Uninfected vs 2003.2_Dissemination | 0.020 | Positive | 1.000 | No |
| 15 | 2003.2_Uninfected vs 2017.1_Uninfected | 0.007 | Positive | 1.000 | No |
| 15 | 2003.2_Uninfected vs 2017.1_Infected | -0.020 | Negative | 1.000 | No |
| 15 | 2003.2_Uninfected vs 2017.1_Disseminated | | | 1.000 | No |
| 15 | 2003.2_Infected vs 2003.2_Dissemination | 0.000 | Negative | 1.000 | No |
| 15 | 2003.2_Infected vs 2017.1_Uninfected | -0.013 | Negative | 1.000 | No |
| 15 | 2003.2_Infected vs 2017.1_Infected | -0.040 | Negative | 1.000 | No |
| 15 | 2003.2_Infected vs 2017.1_Disseminated | | | 1.000 | No |
| 15 | 2003.2_Dissemination vs 2017.1_Uninfected | -0.013 | Negative | 1.000 | No |
| 15 | 2003.2_Dissemination vs 2017.1_Infected | -0.040 | Negative | 1.000 | No |
| 15 | 2003.2_Dissemination vs 2017.1_Disseminated | | | 1.000 | No |
| 15 | 2017.1_Uninfected vs 2017.1_Infected | -0.027 | Negative | 1.000 | No |
| 15 | 2017.1_Uninfected vs 2017.1_Disseminated | | | 1.000 | No |

| | | | | | |
|---|---|---|---|---|---|
| 15 | 2017.1_Infected vs 2017.1_Disseminated | | | 1.000 | No |
| 20 | Unexposed vs 2003.2_Uninfected | 0.289 | Positive | 0.396 | No |
| 20 | Unexposed vs 2003.2_Infected | 0.242 | Positive | 0.002 | Yes |
| 20 | Unexposed vs 2003.2_Dissemination | 0.234 | Positive | 0.010 | Yes |
| 20 | Unexposed vs 2017.1_Uninfected | 0.209 | Positive | 0.745 | No |
| 20 | Unexposed vs 2017.1_Infected | 0.255 | Positive | 0.000 | Yes |
| 20 | Unexposed vs 2017.1_Disseminated | 0.244 | Positive | 0.001 | Yes |
| 20 | 2003.2_Uninfected vs 2003.2_Infected | -0.047 | Negative | 1.000 | No |
| 20 | 2003.2_Uninfected vs 2003.2_Dissemination | -0.055 | Negative | 1.000 | No |
| 20 | 2003.2_Uninfected vs 2017.1_Uninfected | -0.080 | Negative | 1.000 | No |
| 20 | 2003.2_Uninfected vs 2017.1_Infected | -0.034 | Negative | 1.000 | No |
| 20 | 2003.2_Uninfected vs 2017.1_Disseminated | -0.045 | Negative | 1.000 | No |
| 20 | 2003.2_Infected vs 2003.2_Dissemination | -0.008 | Negative | 1.000 | No |
| 20 | 2003.2_Infected vs 2017.1_Uninfected | -0.033 | Negative | 1.000 | No |
| 20 | 2003.2_Infected vs 2017.1_Infected | 0.013 | Positive | 1.000 | No |
| 20 | 2003.2_Infected vs 2017.1_Disseminated | 0.002 | Positive | 1.000 | No |
| 20 | 2003.2_Dissemination vs 2017.1_Uninfected | -0.025 | Negative | 1.000 | No |
| 20 | 2003.2_Dissemination vs 2017.1_Infected | 0.021 | Positive | 1.000 | No |
| 20 | 2003.2_Dissemination vs 2017.1_Disseminated | 0.010 | Positive | 1.000 | No |
| 20 | 2017.1_Uninfected vs 2017.1_Infected | 0.046 | Positive | 1.000 | No |
| 20 | 2017.1_Uninfected vs 2017.1_Disseminated | 0.035 | Positive | 1.000 | No |
| 20 | 2017.1_Infected vs 2017.1_Disseminated | -0.011 | Negative | 1.000 | No |
| 25 | Unexposed vs 2003.2_Uninfected | 0.033 | Positive | 1.000 | No |
| 25 | Unexposed vs 2003.2_Infected | 0.150 | Positive | 0.781 | No |
| 25 | Unexposed vs 2003.2_Dissemination | 0.145 | Positive | 1.000 | No |
| 25 | Unexposed vs 2017.1_Uninfected | 0.250 | Positive | 1.000 | No |
| 25 | Unexposed vs 2017.1_Infected | 0.185 | Positive | 0.252 | No |
| 25 | Unexposed vs 2017.1_Disseminated | 0.180 | Positive | 0.448 | No |
| 25 | 2003.2_Uninfected vs 2003.2_Infected | 0.117 | Positive | 1.000 | No |
| 25 | 2003.2_Uninfected vs 2003.2_Dissemination | 0.112 | Positive | 1.000 | No |
| 25 | 2003.2_Uninfected vs 2017.1_Uninfected | 0.217 | Positive | 1.000 | No |
| 25 | 2003.2_Uninfected vs 2017.1_Infected | 0.152 | Positive | 1.000 | No |
| 25 | 2003.2_Uninfected vs 2017.1_Disseminated | 0.148 | Positive | 1.000 | No |
| 25 | 2003.2_Infected vs 2003.2_Dissemination | -0.005 | Negative | 1.000 | No |
| 25 | 2003.2_Infected vs 2017.1_Uninfected | 0.100 | Positive | 1.000 | No |
| 25 | 2003.2_Infected vs 2017.1_Infected | 0.035 | Positive | 1.000 | No |
| 25 | 2003.2_Infected vs 2017.1_Disseminated | 0.030 | Positive | 1.000 | No |
| 25 | 2003.2_Dissemination vs 2017.1_Uninfected | 0.105 | Positive | 1.000 | No |
| 25 | 2003.2_Dissemination vs 2017.1_Infected | 0.040 | Positive | 1.000 | No |
| 25 | 2003.2_Dissemination vs 2017.1_Disseminated | 0.035 | Positive | 1.000 | No |
| 25 | 2017.1_Uninfected vs 2017.1_Infected | -0.065 | Negative | 1.000 | No |
| 25 | 2017.1_Uninfected vs 2017.1_Disseminated | -0.070 | Negative | 1.000 | No |
| 25 | 2017.1_Infected vs 2017.1_Disseminated | -0.005 | Negative | 1.000 | No |

| | | | | | |
|---|---|---|---|---|---|
| 30 | Unexposed vs 2003.2_Uninfected | -0.055 | Negative | 1.000 | No |
| 30 | Unexposed vs 2003.2_Infected | 0.070 | Positive | 1.000 | No |
| 30 | Unexposed vs 2003.2_Dissemination | 0.070 | Positive | 1.000 | No |
| 30 | Unexposed vs 2017.1_Uninfected | 0.070 | Positive | 1.000 | No |
| 30 | Unexposed vs 2017.1_Infected | -0.015 | Negative | 1.000 | No |
| 30 | Unexposed vs 2017.1_Disseminated | -0.019 | Negative | 1.000 | No |
| 30 | 2003.2_Uninfected vs 2003.2_Infected | 0.125 | Positive | 1.000 | No |
| 30 | 2003.2_Uninfected vs 2003.2_Dissemination | 0.125 | Positive | 1.000 | No |
| 30 | 2003.2_Uninfected vs 2017.1_Uninfected | 0.125 | Positive | 1.000 | No |
| 30 | 2003.2_Uninfected vs 2017.1_Infected | 0.040 | Positive | 1.000 | No |
| 30 | 2003.2_Uninfected vs 2017.1_Disseminated | 0.036 | Positive | 1.000 | No |
| 30 | 2003.2_Infected vs 2003.2_Dissemination | 0.000 | Negative | 1.000 | No |
| 30 | 2003.2_Infected vs 2017.1_Uninfected | 0.000 | Negative | 1.000 | No |
| 30 | 2003.2_Infected vs 2017.1_Infected | -0.085 | Negative | 1.000 | No |
| 30 | 2003.2_Infected vs 2017.1_Disseminated | -0.089 | Negative | 1.000 | No |
| 30 | 2003.2_Dissemination vs 2017.1_Uninfected | 0.000 | Negative | 1.000 | No |
| 30 | 2003.2_Dissemination vs 2017.1_Infected | -0.085 | Negative | 1.000 | No |
| 30 | 2003.2_Dissemination vs 2017.1_Disseminated | -0.089 | Negative | 1.000 | No |
| 30 | 2017.1_Uninfected vs 2017.1_Infected | -0.085 | Negative | 1.000 | No |
| 30 | 2017.1_Uninfected vs 2017.1_Disseminated | -0.089 | Negative | 1.000 | No |
| 30 | 2017.1_Infected vs 2017.1_Disseminated | -0.005 | Negative | 1.000 | No |
| 33 | Unexposed vs 2003.2_Uninfected | 0.000 | Negative | 1.000 | No |
| 33 | Unexposed vs 2003.2_Infected | 0.000 | Negative | 1.000 | No |
| 33 | Unexposed vs 2003.2_Dissemination | 0.000 | Negative | 1.000 | No |
| 33 | Unexposed vs 2017.1_Uninfected | 0.000 | Negative | 1.000 | No |
| 33 | Unexposed vs 2017.1_Infected | 0.000 | Negative | 1.000 | No |
| 33 | Unexposed vs 2017.1_Disseminated | 0.000 | Negative | 1.000 | No |
| 33 | 2003.2_Uninfected vs 2003.2_Infected | 0.000 | Negative | 1.000 | No |
| 33 | 2003.2_Uninfected vs 2003.2_Dissemination | 0.000 | Negative | 1.000 | No |
| 33 | 2003.2_Uninfected vs 2017.1_Uninfected | 0.000 | Negative | 1.000 | No |
| 33 | 2003.2_Uninfected vs 2017.1_Infected | 0.000 | Negative | 1.000 | No |
| 33 | 2003.2_Uninfected vs 2017.1_Disseminated | 0.000 | Negative | 1.000 | No |
| 33 | 2003.2_Infected vs 2003.2_Dissemination | 0.000 | Negative | 1.000 | No |
| 33 | 2003.2_Infected vs 2017.1_Uninfected | 0.000 | Negative | 1.000 | No |
| 33 | 2003.2_Infected vs 2017.1_Infected | 0.000 | Negative | 1.000 | No |
| 33 | 2003.2_Infected vs 2017.1_Disseminated | 0.000 | Negative | 1.000 | No |
| 33 | 2003.2_Dissemination vs 2017.1_Uninfected | 0.000 | Negative | 1.000 | No |
| 33 | 2003.2_Dissemination vs 2017.1_Infected | 0.000 | Negative | 1.000 | No |
| 33 | 2003.2_Dissemination vs 2017.1_Disseminated | 0.000 | Negative | 1.000 | No |
| 33 | 2017.1_Uninfected vs 2017.1_Infected | 0.000 | Negative | 1.000 | No |
| 33 | 2017.1_Uninfected vs 2017.1_Disseminated | 0.000 | Negative | 1.000 | No |
| 33 | 2017.1_Infected vs 2017.1_Disseminated | 0.000 | Negative | 1.000 | No |
| Cycle | Unexposed vs WN03_Uninfected | -0.109 | Negative | 1.000 | No |
| Cycle | Unexposed vs 2003.2_Infected | -0.131 | Negative | 0.714 | No |

| | | | | | |
|---|---|---|---|---|---|
| Cycle | Unexposed vs 2003.2_Dissemination | -0.137 | Negative | 0.657 | No |
| Cycle | Unexposed vs 2017.1_Uninfected | -0.109 | Negative | 1.000 | No |
| Cycle | Unexposed vs 2017.1_Infected | -0.012 | Negative | 1.000 | No |
| Cycle | Unexposed vs 2017.1_Disseminated | -0.034 | Negative | 1.000 | No |
| Cycle | WN03_Uninfected vs 2003.2_Infected | -0.022 | Negative | 1.000 | No |
| Cycle | WN03_Uninfected vs 2003.2_Dissemination | -0.028 | Negative | 1.000 | No |
| Cycle | WN03_Uninfected vs 2017.1_Uninfected | 0.000 | Negative | 1.000 | No |
| Cycle | WN03_Uninfected vs 2017.1_Infected | 0.097 | Positive | 1.000 | No |
| Cycle | WN03_Uninfected vs 2017.1_Disseminated | 0.075 | Positive | 1.000 | No |
| Cycle | 2003.2_Infected vs 2003.2_Dissemination | -0.006 | Negative | 1.000 | No |
| Cycle | 2003.2_Infected vs 2017.1_Uninfected | 0.022 | Positive | 1.000 | No |
| Cycle | 2003.2_Infected vs 2017.1_Infected | 0.120 | Positive | 1.000 | No |
| Cycle | 2003.2_Infected vs 2017.1_Disseminated | 0.097 | Positive | 1.000 | No |
| Cycle | 2003.2_Dissemination vs 2017.1_Uninfected | 0.028 | Positive | 1.000 | No |
| Cycle | 2003.2_Dissemination vs 2017.1_Infected | 0.126 | Positive | 1.000 | No |
| Cycle | 2003.2_Dissemination vs 2017.1_Disseminated | 0.103 | Positive | 1.000 | No |
| Cycle | 2017.1_Uninfected vs 2017.1_Infected | 0.097 | Positive | 1.000 | No |
| Cycle | 2017.1_Uninfected vs 2017.1_Disseminated | 0.075 | Positive | 1.000 | No |
| Cycle | 2017.1_Infected vs 2017.1_Disseminated | -0.022 | Negative | 1.000 | No |

**Table S3. Pairwise differences in ovipositing proportions among infection statuses within each temperature were evaluated using Monte Carlo simulation with Bonferroni correction.**

| Temperature | Pairwise | Magnitude | Direction | P-value | Significance |
|---|---|---|---|---|---|
| 10 | Unexposed - WN02_Uninfected | 0.120 | Negative | 1.000 | No |
| 10 | Unexposed - WN02_Infected | 0.060 | Negative | 1.000 | No |
| 10 | Unexposed - WN02_Disseminated | 0.060 | Negative | 1.000 | No |
| 10 | Unexposed - NY10_Uninfected | 0.000 | Zero | | No |
| 10 | Unexposed - NY10_Infected | 0.000 | Zero | | No |
| 10 | Unexposed - NY10_Disseminated | 0.000 | Zero | | No |
| 10 | WN02_Uninfected - WN02_Infected | 0.060 | Positive | 1.000 | No |
| 10 | WN02_Uninfected - WN02_Disseminated | 0.060 | Positive | 1.000 | No |
| 10 | WN02_Uninfected - NY10_Uninfected | 0.120 | Positive | 0.422 | No |
| 10 | WN02_Uninfected - NY10_Infected | 0.120 | Positive | 0.407 | No |
| 10 | WN02_Uninfected - NY10_Disseminated | 0.120 | Positive | 0.402 | No |
| 10 | WN02_Infected - WN02_Disseminated | 0.000 | Zero | 1.000 | No |
| 10 | WN02_Infected - NY10_Uninfected | 0.060 | Positive | 1.000 | No |
| 10 | WN02_Infected - NY10_Infected | 0.060 | Positive | 1.000 | No |
| 10 | WN02_Infected - NY10_Disseminated | 0.060 | Positive | 1.000 | No |
| 10 | WN02_Disseminated - NY10_Uninfected | 0.060 | Positive | 1.000 | No |
| 10 | WN02_Disseminated - NY10_Infected | 0.060 | Positive | 1.000 | No |
| 10 | WN02_Disseminated - NY10_Disseminated | 0.060 | Positive | 1.000 | No |
| 10 | NY10_Uninfected - NY10_Infected | 0.000 | Zero | | No |
| 10 | NY10_Uninfected - NY10_Disseminated | 0.000 | Zero | | No |
| 10 | NY10_Infected - NY10_Disseminated | 0.000 | Zero | | No |
| 15 | Unexposed - WN02_Uninfected | 0.493 | Positive | 0.000 | Yes |
| 15 | Unexposed - WN02_Infected | 0.580 | Positive | 0.000 | Yes |
| 15 | Unexposed - WN02_Disseminated | 0.601 | Positive | 0.000 | Yes |
| 15 | Unexposed - NY10_Uninfected | 0.422 | Positive | 0.008 | Yes |
| 15 | Unexposed - NY10_Infected | 0.565 | Positive | 0.000 | Yes |
| 15 | Unexposed - NY10_Disseminated | 0.667 | Positive | 0.000 | Yes |
| 15 | WN02_Uninfected - WN02_Infected | 0.087 | Positive | 1.000 | No |
| 15 | WN02_Uninfected - WN02_Disseminated | 0.109 | Positive | 1.000 | No |
| 15 | WN02_Uninfected - NY10_Uninfected | 0.071 | Negative | 1.000 | No |
| 15 | WN02_Uninfected - NY10_Infected | 0.072 | Positive | 1.000 | No |
| 15 | WN02_Uninfected - NY10_Disseminated | 0.174 | Positive | 0.052 | No |
| 15 | WN02_Infected - WN02_Disseminated | 0.022 | Positive | 1.000 | No |
| 15 | WN02_Infected - NY10_Uninfected | 0.158 | Negative | 1.000 | No |
| 15 | WN02_Infected - NY10_Infected | 0.015 | Negative | 1.000 | No |
| 15 | WN02_Infected - NY10_Disseminated | 0.087 | Positive | 1.000 | No |
| 15 | WN02_Disseminated - NY10_Uninfected | 0.180 | Negative | 0.466 | No |
| 15 | WN02_Disseminated - NY10_Infected | 0.037 | Negative | 1.000 | No |
| 15 | WN02_Disseminated - NY10_Disseminated | 0.065 | Positive | 1.000 | No |
| 15 | NY10_Uninfected - NY10_Infected | 0.143 | Positive | 1.000 | No |
| 15 | NY10_Uninfected - NY10_Disseminated | 0.245 | Positive | 0.007 | Yes |

| | | | | | |
|---|---|---|---|---|---|
| 15 | NY10_Infected - NY10_Disseminated | 0.102 | Positive | 1.000 | No |
| 20 | Unexposed - WN02_Uninfected | 0.618 | Positive | 0.000 | Yes |
| 20 | Unexposed - WN02_Infected | 0.292 | Positive | 0.425 | No |
| 20 | Unexposed - WN02_Disseminated | 0.394 | Positive | 0.021 | Yes |
| 20 | Unexposed - NY10_Uninfected | 0.598 | Positive | 0.000 | Yes |
| 20 | Unexposed - NY10_Infected | 0.210 | Positive | 1.000 | No |
| 20 | Unexposed - NY10_Disseminated | 0.373 | Positive | 0.049 | Yes |
| 20 | WN02_Uninfected - WN02_Infected | 0.327 | Negative | 0.006 | Yes |
| 20 | WN02_Uninfected - WN02_Disseminated | 0.224 | Negative | 0.194 | No |
| 20 | WN02_Uninfected - NY10_Uninfected | 0.020 | Negative | 1.000 | No |
| 20 | WN02_Uninfected - NY10_Infected | 0.408 | Negative | 0.000 | Yes |
| 20 | WN02_Uninfected - NY10_Disseminated | 0.245 | Negative | 0.111 | No |
| 20 | WN02_Infected - WN02_Disseminated | 0.102 | Positive | 1.000 | No |
| 20 | WN02_Infected - NY10_Uninfected | 0.306 | Positive | 0.020 | Yes |
| 20 | WN02_Infected - NY10_Infected | 0.082 | Negative | 1.000 | No |
| 20 | WN02_Infected - NY10_Disseminated | 0.082 | Positive | 1.000 | No |
| 20 | WN02_Disseminated - NY10_Uninfected | 0.204 | Positive | 0.487 | No |
| 20 | WN02_Disseminated - NY10_Infected | 0.184 | Negative | 1.000 | No |
| 20 | WN02_Disseminated - NY10_Disseminated | 0.020 | Negative | 1.000 | No |
| 20 | NY10_Uninfected - NY10_Infected | 0.388 | Negative | 0.001 | Yes |
| 20 | NY10_Uninfected - NY10_Disseminated | 0.224 | Negative | 0.261 | No |
| 20 | NY10_Infected - NY10_Disseminated | 0.163 | Positive | 1.000 | No |
| 25 | Unexposed - WN02_Uninfected | 0.596 | Positive | 0.000 | Yes |
| 25 | Unexposed - WN02_Infected | 0.290 | Positive | 0.331 | No |
| 25 | Unexposed - WN02_Disseminated | 0.351 | Positive | 0.049 | Yes |
| 25 | Unexposed - NY10_Uninfected | 0.733 | Positive | 0.000 | Yes |
| 25 | Unexposed - NY10_Infected | 0.156 | Positive | 1.000 | No |
| 25 | Unexposed - NY10_Disseminated | 0.333 | Positive | 0.159 | No |
| 25 | WN02_Uninfected - WN02_Infected | 0.306 | Negative | 0.065 | No |
| 25 | WN02_Uninfected - WN02_Disseminated | 0.245 | Negative | 0.372 | No |
| 25 | WN02_Uninfected - NY10_Uninfected | 0.137 | Positive | 1.000 | No |
| 25 | WN02_Uninfected - NY10_Infected | 0.440 | Negative | 0.000 | Yes |
| 25 | WN02_Uninfected - NY10_Disseminated | 0.263 | Negative | 0.178 | No |
| 25 | WN02_Infected - WN02_Disseminated | 0.061 | Positive | 1.000 | No |
| 25 | WN02_Infected - NY10_Uninfected | 0.444 | Positive | 0.000 | Yes |
| 25 | WN02_Infected - NY10_Infected | 0.134 | Negative | 1.000 | No |
| 25 | WN02_Infected - NY10_Disseminated | 0.044 | Positive | 1.000 | No |
| 25 | WN02_Disseminated - NY10_Uninfected | 0.382 | Positive | 0.001 | Yes |
| 25 | WN02_Disseminated - NY10_Infected | 0.195 | Negative | 1.000 | No |
| 25 | WN02_Disseminated - NY10_Disseminated | 0.018 | Negative | 1.000 | No |
| 25 | NY10_Uninfected - NY10_Infected | 0.578 | Negative | 0.000 | Yes |
| 25 | NY10_Uninfected - NY10_Disseminated | 0.400 | Negative | 0.000 | Yes |
| 25 | NY10_Infected - NY10_Disseminated | 0.178 | Positive | 1.000 | No |
| 30 | Unexposed - WN02_Uninfected | 0.439 | Positive | 0.002 | Yes |

| | | | | | |
|---|---|---|---|---|---|
| 30 | Unexposed - WN02_Infected | 0.205 | Positive | 1.000 | No |
| 30 | Unexposed - WN02_Disseminated | 0.269 | Positive | 0.642 | No |
| 30 | Unexposed - NY10_Uninfected | 0.507 | Positive | 0.000 | Yes |
| 30 | Unexposed - NY10_Infected | 0.007 | Positive | 1.000 | No |
| 30 | Unexposed - NY10_Disseminated | 0.027 | Positive | 1.000 | No |
| 30 | WN02_Uninfected - WN02_Infected | 0.234 | Negative | 0.327 | No |
| 30 | WN02_Uninfected - WN02_Disseminated | 0.170 | Negative | 1.000 | No |
| 30 | WN02_Uninfected - NY10_Uninfected | 0.068 | Positive | 1.000 | No |
| 30 | WN02_Uninfected - NY10_Infected | 0.432 | Negative | 0.001 | Yes |
| 30 | WN02_Uninfected - NY10_Disseminated | 0.412 | Negative | 0.001 | Yes |
| 30 | WN02_Infected - WN02_Disseminated | 0.064 | Positive | 1.000 | No |
| 30 | WN02_Infected - NY10_Uninfected | 0.302 | Positive | 0.005 | Yes |
| 30 | WN02_Infected - NY10_Infected | 0.198 | Negative | 1.000 | No |
| 30 | WN02_Infected - NY10_Disseminated | 0.178 | Negative | 1.000 | No |
| 30 | WN02_Disseminated - NY10_Uninfected | 0.238 | Positive | 0.059 | No |
| 30 | WN02_Disseminated - NY10_Infected | 0.262 | Negative | 0.278 | No |
| 30 | WN02_Disseminated - NY10_Disseminated | 0.242 | Negative | 0.480 | No |
| 30 | NY10_Uninfected - NY10_Infected | 0.500 | Negative | 0.000 | Yes |
| 30 | NY10_Uninfected - NY10_Disseminated | 0.480 | Negative | 0.000 | Yes |
| 30 | NY10_Infected - NY10_Disseminated | 0.020 | Positive | 1.000 | No |
| 33 | Unexposed - WN02_Uninfected | 0.133 | Positive | 0.288 | No |
| 33 | Unexposed - WN02_Infected | 0.133 | Positive | 0.281 | No |
| 33 | Unexposed - WN02_Disseminated | 0.133 | Positive | 0.289 | No |
| 33 | Unexposed - NY10_Uninfected | 0.133 | Positive | 0.331 | No |
| 33 | Unexposed - NY10_Infected | 0.068 | Positive | 1.000 | No |
| 33 | Unexposed - NY10_Disseminated | 0.068 | Positive | 1.000 | No |
| 33 | WN02_Uninfected - WN02_Infected | 0.000 | Zero | | No |
| 33 | WN02_Uninfected - WN02_Disseminated | 0.000 | Zero | | No |
| 33 | WN02_Uninfected - NY10_Uninfected | 0.000 | Zero | | No |
| 33 | WN02_Uninfected - NY10_Infected | 0.065 | Negative | 1.000 | No |
| 33 | WN02_Uninfected - NY10_Disseminated | 0.065 | Negative | 1.000 | No |
| 33 | WN02_Infected - WN02_Disseminated | 0.000 | Zero | | No |
| 33 | WN02_Infected - NY10_Uninfected | 0.000 | Zero | | No |
| 33 | WN02_Infected - NY10_Infected | 0.065 | Negative | 1.000 | No |
| 33 | WN02_Infected - NY10_Disseminated | 0.065 | Negative | 1.000 | No |
| 33 | WN02_Disseminated - NY10_Uninfected | 0.000 | Zero | | No |
| 33 | WN02_Disseminated - NY10_Infected | 0.065 | Negative | 1.000 | No |
| 33 | WN02_Disseminated - NY10_Disseminated | 0.065 | Negative | 1.000 | No |
| 33 | NY10_Uninfected - NY10_Infected | 0.065 | Negative | 1.000 | No |
| 33 | NY10_Uninfected - NY10_Disseminated | 0.065 | Negative | 1.000 | No |
| 33 | NY10_Infected - NY10_Disseminated | 0.000 | Zero | 1.000 | No |
| Cycle | Unexposed - WN02_Uninfected | 0.724 | Positive | 0.000 | Yes |
| Cycle | Unexposed - WN02_Infected | 0.192 | Positive | 1.000 | No |
| Cycle | Unexposed - WN02_Disseminated | 0.235 | Positive | 1.000 | No |
| Cycle | Unexposed - NY10_Uninfected | 0.663 | Positive | 0.000 | Yes |

| | | | | | |
|---|---|---|---|---|---|
| Cycle | Unexposed - NY10_Infected | 0.288 | Positive | 0.384 | No |
| Cycle | Unexposed - NY10_Disseminated | 0.392 | Positive | 0.023 | Yes |
| Cycle | WN02_Uninfected - WN02_Infected | 0.532 | Negative | 0.000 | Yes |
| Cycle | WN02_Uninfected - WN02_Disseminated | 0.489 | Negative | 0.000 | Yes |
| Cycle | WN02_Uninfected - NY10_Uninfected | 0.062 | Negative | 1.000 | No |
| Cycle | WN02_Uninfected - NY10_Infected | 0.437 | Negative | 0.000 | Yes |
| Cycle | WN02_Uninfected - NY10_Disseminated | 0.332 | Negative | 0.002 | Yes |
| Cycle | WN02_Infected - WN02_Disseminated | 0.043 | Positive | 1.000 | No |
| Cycle | WN02_Infected - NY10_Uninfected | 0.470 | Positive | 0.000 | Yes |
| Cycle | WN02_Infected - NY10_Infected | 0.095 | Positive | 1.000 | No |
| Cycle | WN02_Infected - NY10_Disseminated | 0.199 | Positive | 1.000 | No |
| Cycle | WN02_Disseminated - NY10_Uninfected | 0.428 | Positive | 0.000 | Yes |
| Cycle | WN02_Disseminated - NY10_Infected | 0.053 | Positive | 1.000 | No |
| Cycle | WN02_Disseminated - NY10_Disseminated | 0.157 | Positive | 1.000 | No |
| Cycle | NY10_Uninfected - NY10_Infected | 0.375 | Negative | 0.002 | Yes |
| Cycle | NY10_Uninfected - NY10_Disseminated | 0.271 | Negative | 0.076 | No |
| Cycle | NY10_Infected - NY10_Disseminated | 0.104 | Positive | 1.000 | No |
